# Transcript expression-aware annotation improves rare variant discovery and interpretation

**DOI:** 10.1101/554444

**Authors:** Beryl B. Cummings, Konrad J. Karczewski, Jack A. Kosmicki, Eleanor G. Seaby, Nicholas A. Watts, Moriel Singer-Berk, Jonathan M. Mudge, Juha Karjalainen, F. Kyle Satterstrom, Anne O’Donnell-Luria, Timothy Poterba, Cotton Seed, Matthew Solomonson, Jessica Alföldi, The Genome Aggregation Database Production Team, The Genome Aggregation Database Consortium, Mark J. Daly, Daniel G. MacArthur

## Abstract

The acceleration of DNA sequencing in patients and population samples has resulted in unprecedented catalogues of human genetic variation, but the interpretation of rare genetic variants discovered using such technologies remains extremely challenging. A striking example of this challenge is the existence of disruptive variants in dosage-sensitive disease genes, even in apparently healthy individuals. Through manual curation of putative loss of function (pLoF) variants in haploinsufficient disease genes in the Genome Aggregation Database (gnomAD)(*1*), we show that one explanation for this paradox involves alternative mRNA splicing, which allows exons of a gene to be expressed at varying levels across cell types. Currently, no existing annotation tool systematically incorporates this exon expression information into variant interpretation. Here, we develop a transcript-level annotation metric, the proportion expressed across transcripts (pext), which summarizes isoform quantifications for variants. We calculate this metric using 11,706 tissue samples from the Genotype Tissue Expression project(*2*) (GTEx) and show that it clearly differentiates between weakly and highly evolutionarily conserved exons, a proxy for functional importance. We demonstrate that expression-based annotation selectively filters 22.8% of falsely annotated pLoF variants found in haploinsufficient disease genes in gnomAD, while removing less than 4% of high-confidence pathogenic variants in the same genes. Finally, we apply our expression filter to the analysis of *de novo* variants in patients with autism spectrum disorder (ASD) and developmental disorders and intellectual disability (DD/ID) to show that pLoF variants in weakly expressed regions have effect sizes similar to those of synonymous variants, while pLoF variants in highly expressed exons are most strongly enriched among cases versus controls. Our annotation is fast, flexible, and generalizable, making it possible for any variant file to be annotated with any isoform expression dataset, and will be valuable for rare disease diagnosis, rare variant burden analyses in complex disorders, and curation and prioritization of variants in recall-by-genotype studies.

---

A primary challenge in the use of genome and exome sequencing to predict human phenotypes is that our capacity to identify genetic variation exceeds our ability to interpret their functional impact (*3, 4*). One underappreciated source of variability for variant interpretation involves differences in alternative mRNA splicing, which enables exons to be expressed at different levels across tissues. These expression differences mean that variants in different regions of a gene can have different phenotypic outcomes depending on the isoforms they affect. For example, variants occurring in an exon differentially included in two isoforms of *CACNA1C* with diverse tissue expression patterns result in distinct types of Timothy syndrome (*5*). Pathogenic variants in the isoform that exhibits multi-tissue expression result in a multi-system disorder (*5-7*), whereas those on the isoform predominantly expressed in heart result in more severe and specific cardiac defects (*8*). In addition, Mendelian variants have been found on tissue-specific isoforms (*9, 10*) and isoform expression levels in *TTN* have been used to show that pLoF variants found in healthy controls occur in exons that are absent from dominantly expressed isoforms, whereas those in dilated cardiomyopathy patients occur on constitutive exons (*11*), emphasizing the utility of exon expression information for variant interpretation.

We find that isoform diversity is also a contributor to the paradoxical finding of disruptive variants in dosage-sensitive disease genes in ostensibly healthy individuals. In the gnomAD database, we identify 401 high-quality pLoF variants that pass both sequencing and annotation quality filters in 61 haploinsufficient disease genes where heterozygous pLoF variants are established to cause severe developmental delay phenotypes with high penetrance (Methods). Given the severity of these phenotypes and their extremely low worldwide prevalence, ranging from 1 in 10,000 to less than 1 in a million, very few, if any true pLoF variants would be expected to be found in the gnomAD population. As such, most or all of these observed pLoF variants are likely to be sequencing or annotation errors (*12*). Manual curation of these variants reveals common error modes that result in likely misannotation of pLoFs, with diversity of transcript structure, mediated by variants falling on low-confidence transcripts, emerging as a major consideration (Figure 1, Supplementary Figure 1, Supplementary Tables 1-3). However, no existing tools systematically incorporate information on transcript expression into variant interpretation.

**Figure 1:**
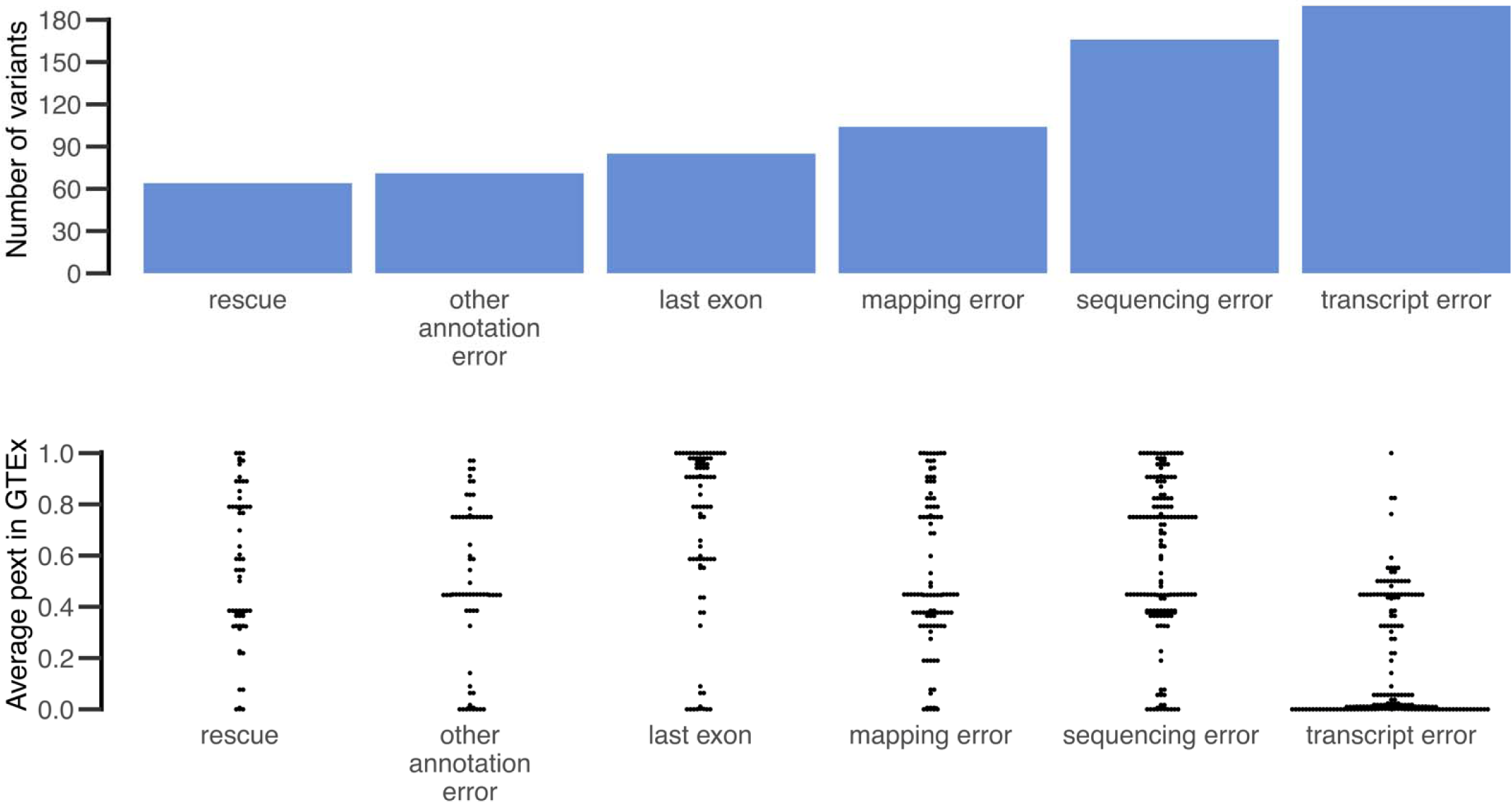
Curation of pLoF variants in haploinsufficient disease genes found in gnomAD reveals transcript errors as a major confounding error mode in variant annotation. We identified and manually curated 401 pLoF variants in the gnomAD dataset in 61 haploinsufficient severe developmental delay genes and flagged any reason the pLoF may not be a true LoF variant. Top plot shows the frequency of each error mode present in the 306 variants classified as unlikely to be a true LoF. Transcript errors emerge as a major putative error mode in the annotation of these pLoF variants. Beeswarm plot on bottom shows the average pext score across GTEx tissues presented in the manuscript for each variant in the error categories. This shows that pext values are discriminately lower for variants that are annotated as possible transcript errors.

The advent of large-scale transcriptome sequencing datasets, such as GTEx (*2*), provides an opportunity to incorporate cross-tissue exon expression into variant interpretation. However, the current formats of these databases do not readily allow for unbiased estimation of exon expression. The GTEx web browser offers information on exon-level read pileup across tissues, but this approach is confounded by technical artifacts such as 3’ bias (*13*) (preferential coverage of bases close to the 3’ end of a transcript; Supplementary Figure 2A). Such systematic biases mean that simple exon-level coverage in a transcriptome dataset cannot be used as a reliable proxy for exon expression, especially in longer genes (Figure 2A, Supplementary Figure 2B).

**Figure 2:**
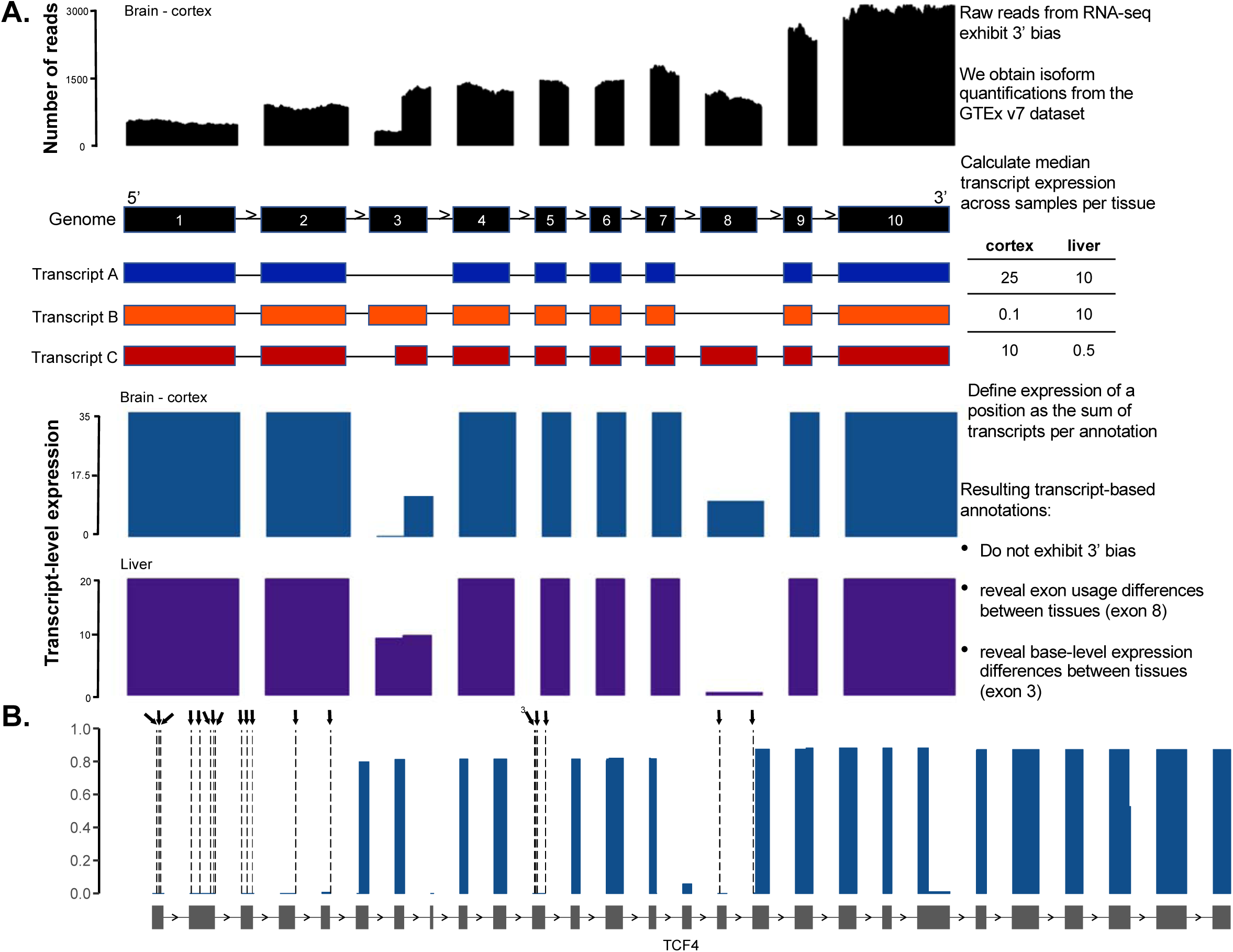
Summary of transcript-expression based annotation method. **A.** Overview of transcript aware annotation. Most genes have many annotated isoforms, which can have varying expression patterns across tissues. Utilizing number of reads aligning to exonic regions in transcriptome datasets as a proxy for exon expression (top panel black) has confounding effects due to 3’ bias. In this example, while exon 3 and 8 may have different expression levels in Brain – Cortex, the number of reads aligning to the two exons are similar, masking exon usage differences. Transcript-aware annotation defines the expression of every variant as the sum of transcripts that have the same annotation. The resulting transcript-level expression plots do not exhibit 3’ bias, and reveal exon usage differences across tissues. **B.** Example of utility of transcript-expression based annotation. There are 20 high quality pLoF variants in the haploinsufficient developmental delay gene *TCF4* in gnomAD, annotated as dashed lines and arrows. All 20 variants have no evidence of expression in the GTEx dataset, suggesting functional TCF4 protein can be made in the presence of these variants.

Isoform quantification tools provide estimates of isoform expression levels that correct, albeit imperfectly(*13, 14*), for confounding by 3’ bias as well as other technical artifacts such as isoform length, isoform GC content, and transcript sequence complexity (*15-17*). Here, we utilize isoform-level quantifications from 11,706 tissue samples from the GTEx v7 dataset to derive an annotation-specific expression metric. For each tissue, we annotate each variant with the expression of every possible consequence across all transcripts, which can be used to summarize expression in any combination of tissues of interest. We first compute the median expression of a transcript across tissue samples, and define the expression of a given variant as the sum of the expression of all transcripts for which the variant has the same annotation (Figure 2A, Supplementary the expression of the annotation to the total gene expression, we define a metric (proportion expression across transcripts, or pext), which can be interpreted as a measure of the proportion of the total transcriptional output from a gene that would be affected by the variant annotation in question (Supplementary Figure 3B).

**Figure 3:**
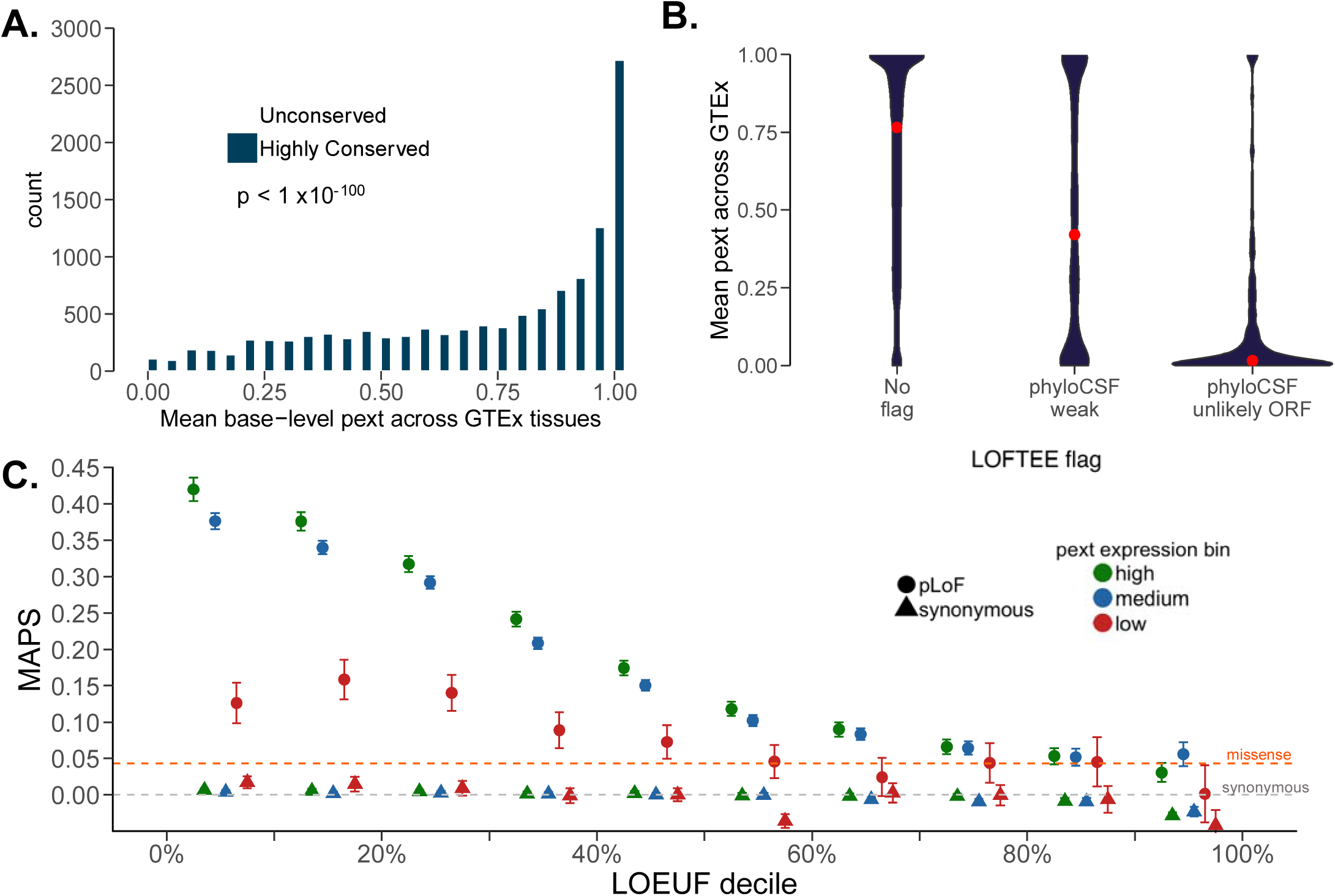
Functional validation of transcript-expression based annotation. **A.** We define highly conserved and unconserved regions and compared the expression status of these regions across GTEx. Highly conserved regions are enriched for having near-constitutive expression whereas unconserved regions are enriched for having little to no usage across GTEx. This difference is significant after correcting for gene length (logistic regression p value < 1 × 10^−100^). We note that unconserved regions with high levels of expression (pext > 0.9) are enriched for immune-related genes, which are selected for diversity and thus have low conservation, but represent true coding regions. **B.** Transcript-expression based annotation recapitulates, and adds information to, existing interpretation tools. LOFTEE-HC pLoF variants in gnomAD with no flags are enriched for higher pext values, whereas HC pLoF variants falling on low phyloCSF or unlikely ORF regions are enriched for low expression. However, HC-pLoF variants can also be filtered based on a low pext score. Red dots represent median pext value across GTEx tissues. **C.** Nonsynonymous variants found on near-constitutive regions tend to be more deleterious. We compared the mutability adjusted proportion singleton (MAPS) score for variants with low (<0.1), medium (0.1 ≤ pext ≤ 0.9) and high (pext > 0.9) expression. Variants with near-constitutive expression have a higher MAPS score, indicating higher deleteriousness than those with little to no evidence of expression. Dashed grey and orange line represent MAPS values for all gnomAD missense and all synonymous variants, respectively.

The pext metric allows for quick visualization of the expression of exons across a gene. Figure 2B shows *TCF4*, a haploinsufficient gene in which heterozygous variants result in Pitt-Hopkins syndrome (*18*), a highly penetrant disorder associated with severe developmental delay. This gene harbors 20 unique high quality pLoF mutations across 56 individuals in the gnomAD database. All 20 variants lie on exons with no evidence of expression across the GTEx dataset (Figure 2B, Supplementary Figure 4) indicating that functional TCF4 protein can be made in the presence of these variants. This visualization is now available for all genes in the gnomAD browser (gnomad.broadinstitute.org), and can aid in rapid identification of variants occurring on exons with little to no evidence of expression in GTEx.

**Figure 4:**
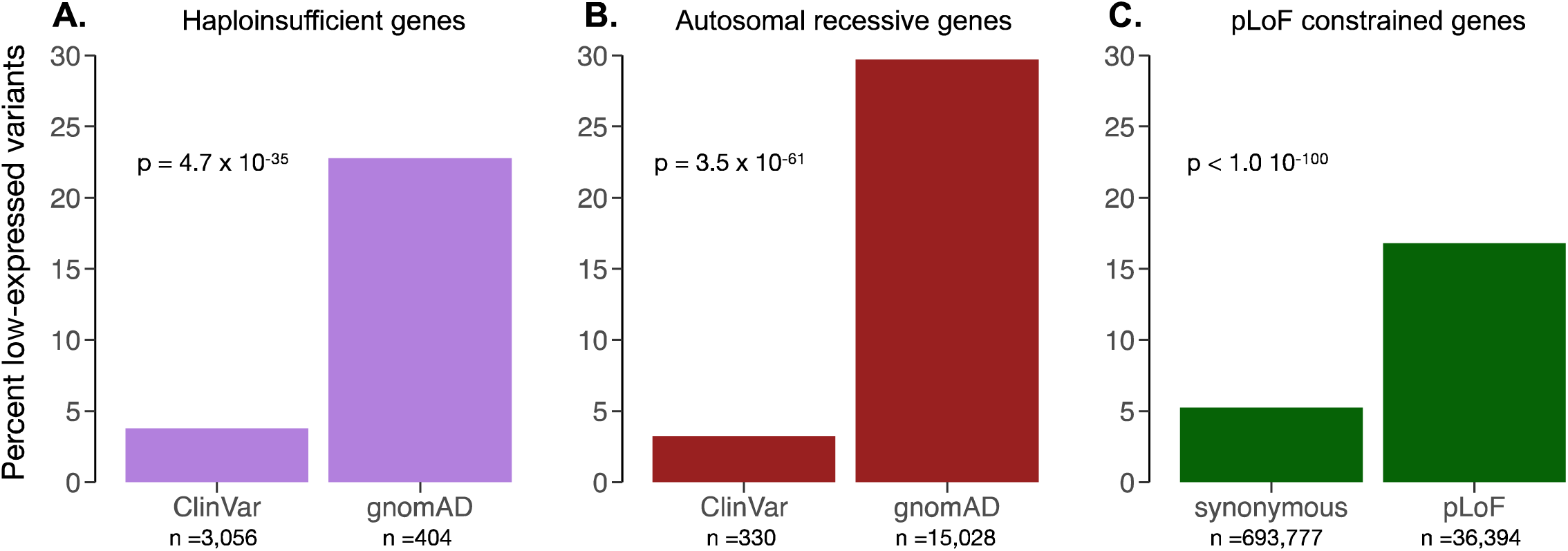
Transcript-expression based annotation aids Mendelian variant interpretation. **A.** Comparison of proportion of high quality pLoF variants filtered in a curated list of 61 haploinsufficient developmental delays genes in gnomAD vs ClinVar with a cutoff of average pext across GTEx ≤ 0.1 (low expression). Expression-based filtering results in removal of 22.8% of gnomAD pLoFs and 3.8% of confidently curated set of pLoFs in ClinVar. **B.** Expression-based annotation filters 30% of pLoF variants found in gnomAD in a homozygous state in at least one individual, and 3.2% of any pLoF variants found in the same genes in ClinVar. **C.** We extended this filtering approach to pLoF and synonymous variants in gnomAD pLoF-intolerant genes (defined by LOEF < 0.35). This filters 16.8% LoF and 5.2% of synonymous variants. Numbers below bar plots indicate the total number of high-quality variants considered in each group. For pLoFs only LOFTEE-HC variants were considered, p-values calculated from fisher’s exact test for counts.

To explore whether expression-based annotation marks functionally important regions, we compared the distribution of the pext metric in evolutionarily conserved and unconserved regions using phyloCSF (*19*). Exons with patterns of multi-species conservation consistent with coding regions have higher phyloCSF scores, and should exhibit detectable expression patterns, whereas regions with lower scores will be enriched for incorrect exon annotations, which are expected to have little evidence of expression in a population transcriptome dataset. As expected, we observe significantly lower expression for unconserved regions, and near-constitutive expression in highly conserved regions (Figure 3A, Supplementary Figure 5A). This difference remains statistically significant after correcting for exon length (logistic regression p < 1.0 × 10^−100^), which can influence both phyloCSF scores and isoform quantifications, indicating that transcript expression-aware annotation marks functionally relevant exonic regions.

**Figure 5:**
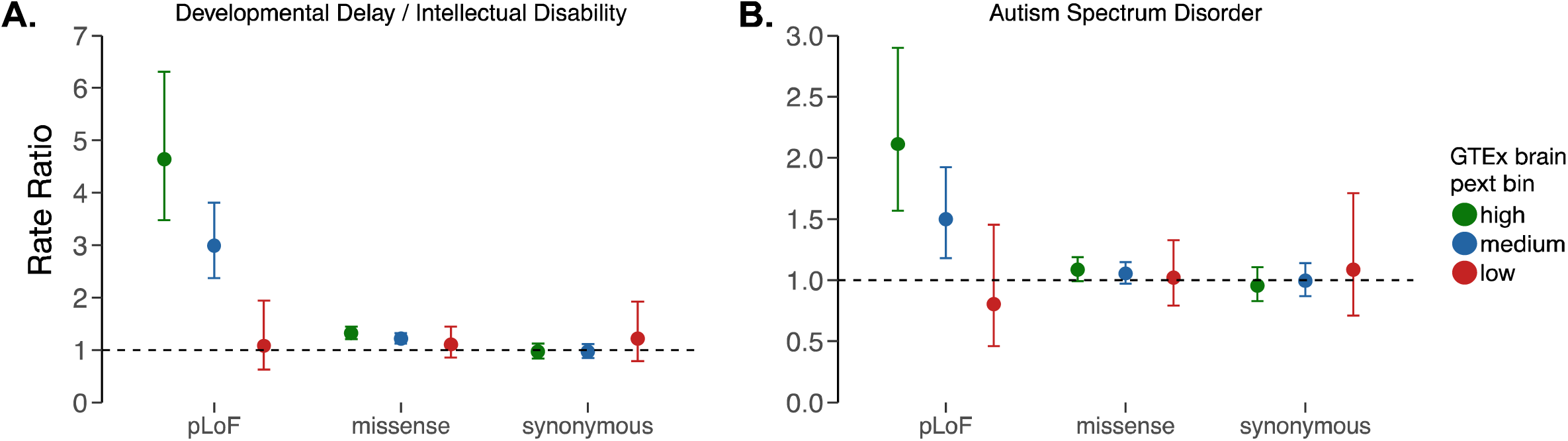
Application of transcript-expression based annotation to *de novo* variant analyses in **A.** developmental delay and/or intellectual disability (DD/ID) and **B.** autism spectrum disorder (ASD). We find that de novo pLoF variants found on near consitutively expressed regions in GTEx brain tissues have larger effect sizes than *de novo* LoF variants in weakly expressed regions in both disorders. Strikingly, *de novo* pLoF variants found on regions with little evidence for expression are equally distributed in cases vs controls as *de novo* synonymous variants, suggesting such variants can be removed from gene burden testing analyses to boost discovery power. The high pext expression bin contains 46.1% and 42.3%, and the low expression bin contains 4.0% and 6.0% of pLoF variants found in DD/ID and ASD patients, respectively. Rate ratio represents estimate from the poisson exact test.

While the metrics are associated, we find that pext provides orthogonal information to conservation for variant interpretation. For example, regions with low evidence of conservation but high expression (in Figure 3A) are enriched for genes in immune-related pathways (Methods), which are selected for diversity but represent true coding regions. In addition, the pext value is higher for pLoF variants annotated as high confidence (HC) by the Loss of Function Transcript Effect Estimator(*1*) (LOFTEE) with no additional flags than those flagged as having found on unlikely open reading frames or weakly conserved regions (Figure 3B, Supplementary Figure 5B). However, LOFTEE-HC variants with no flags can also have low pext values, suggesting transcript-expression aware annotation adds additional information to the currently available interpretation toolkit.

We undertook manual evaluation of 128 regions marked as unexpressed (mean pext < 0.1 in all tissues and in GTEx brain) in 61 haploinsufficient genes following the GENCODE manual annotation workflow(*20*) to evaluate the annotation quality in these coding sequence (CDS) regions. A third of flagged regions were associated with low quality models that have been removed or switched to non-coding biotypes in subsequent GENCODE releases (Supplementary Figure 6) while 70% of the remaining regions correspond to models that satisfy only minimum criteria for inclusion in the gene set, corresponding to ‘putative’ annotations that lack markers for CDS functionality (Supplementary Table 4). Nonetheless, we find support for some highly conserved CDS regions, several of which show evidence of transcription in fetal tissues, underlining the importance of incorporating multiple isoform expression datasets for interpretation (Supplementary Figure 6D).

Nonsynonymous variants found on constitutively expressed regions would be expected to be more deleterious than those on regions with no evidence of expression. To test this, we defined expression bins based on the average pext value across GTEx tissues where an average pext value less than 0.1 was defined as low (or unexpressed), above 0.9 as high (or near-constitutive) and intermediate values as medium expression. We compared the mutability-adjusted proportion singleton (MAPS), a measure of negative selection on variant classes(*21*), partitioned on the LoF Observed Upper-bound Fraction (LOEUF) decile, a measure of constraint against pLoF variants in the gnomAD dataset(*1*) in each of these expression bins. MAPS scores differed substantially between pLoF variants found on low-expressed and high-expressed regions in genes intolerant to pLoF variation (Figure 3C, Supplementary Figure 5C). This information is complementary to existing variant prioritization tools such as PolyPhen-2(*22*) (Supplementary Figure 5D). This skew of nonsynonymous variation in high-expressed regions suggests that variation arising in such exons tends be more deleterious, whereas nonsynonymous variants on regions with low expression are similar to missense variants in their inferred deleteriousness.

To evaluate the utility of transcript expression-based annotation in Mendelian variant interpretation, we assessed the number of variants that would be filtered based on a pext cutoff of <0.1 (low expression) across GTEx tissues for three gene sets. Firstly, we evaluated high-quality pLoF variants in the 61 manually curated haploinsufficient genes in gnomAD and ClinVar(*23*). The low pext expression bin resulted in filtering of 22.8% of pLoF variants in haploinsufficient developmental delay genes in gnomAD, but only 3.8% of high-quality pathogenic variants in ClinVar (Figure 4A; p = 4.7 × 10^−35^ Methods). We next compared pLoF variants in autosomal recessive disease genes found in a homozygous state in at least one individual in gnomAD and any pLoF variant in these genes in ClinVar and observed similar results: expression-based annotation filters 30.0% of variants in gnomAD while only filtering 3.2% of variants in ClinVar (Figure 4B; p = 3.5 × 10^−61^).

Finally, we evaluated gnomAD pLoF variants in genes that are constrained against pLoF variation(*1*) (LOEUF score < 0.35). Given that these genes are depleted for loss-of-function variation in the general population, we expect the observed pLoF variants in these genes to be enriched for annotation errors. We compared the proportion filtered to synonymous variants in the same genes, which we expect to be randomly distributed. Our metric removes 16.8% of pLoF variants in constrained genes, but only 5.2% of synonymous variants (Figure 4C; p < 1.0 × 10^−100^). In all cases, the vast majority of filtered variants were otherwise high-confidence with no LOFTEE annotation flags, suggesting again that pext provided additional information to existing variant prioritization tools in removing annotation errors (Supplementary Figure 7).

To explore the benefits of this approach for rare variant analyses, we applied pext binning to burden testing of *de novo* variants in patients with developmental delay/intellectual disability (DD/ID) or autism spectrum disorder (ASD) using a set of 23,970 *de novo* variants collated from several studies including the Deciphering Developmental Disorders (DDD) project and the Autism Sequencing Consortium (ASC)(*24-29*). We find that *de novo* pLoF variants in patients with DD/ID in low-expressed regions have effect sizes similar to those of synonymous variants (rate ratio, denoted as RR, of low-expressed pLoFs = 1.08, p = 0.90) whereas pLoF variants in highly expressed regions have much larger effect sizes (RR = 4.64, p = 3.74 × 10^−38^; Figure 5A). This observation is consistent for *de novo* variants in autism (RR for low-expressed pLoFs = 0.80, p = 0.47; RR for high-expressed pLoFs = 2.11, p = 8.2 × 10^−8^, Figure 5B) and congenital heart disease with co-morbid neurodevelopmental delay (Supplementary Figure 8A) as well as rare variants (AC ≤ 10) identified in highly constrained genes in the large iPSYCH case/control study of Danish patients with autism spectrum disorder and attention-deficit/hyperactivity disorder (Supplementary Figure 8B). Overall, we consistently observe low-expressed pLoFs to have effect sizes similar to those of synonymous variants, with pLoF variants in constitutive regions having larger effect sizes, suggesting that incorporating transcript expression-aware annotation in rare variant studies can boost power for gene discovery.

We have described the development and validation of a transcript expression-based annotation framework to integrate results from transcriptome sequencing experiments into clinical variant interpretation. While our initial analysis utilizes GTEx, our method can be used with any isoform expression dataset to annotate any variant file rapidly in the scalable software framework Hail (https://hail.is). For example, annotation of >120,000 gnomAD individuals with GTEx takes under an hour using 60 cores, at a cost of about $5 on public cloud compute, which can be further scaled to larger datasets. In addition, the annotations we provide are flexible: while we have described the use of average transcript-level expression across many tissues, alternative approaches such as using maximum expression across any tissue, may prove useful depending on variant interpretation goals (Supplementary Figures 9 & 10).

We note that while this metric successfully discriminates between near-constitutive and low expression levels, which are useful for prioritizing and filtering variants, respectively, regions with intermediate expression levels are more challenging to interpret. However, we hypothesize directed analyses of intermediate expression levels may help elucidate the role of alternative splicing in phenotypic diversity (*30, 31*). In addition, while we have binned average pext scores across GTEx tissues into low, medium and high expression, different genes will likely have varying optimal tissues and thresholds for variant interpretation. Regions tagged as low expression are often corroborated by expert opinion of CDS curation, but domain knowledge of a gene will outperform this summary metric.

An important caveat in our approach is the imprecision of isoform quantification methods using short-read transcriptome data. However, we note that repeating key analyses in the manuscript with a different isoform quantification tool showed consistent results (Methods, Supplementary Figure 11), suggesting robustness to the precise pipeline used. The utility of this framework will increase as our ability to quantify isoform expression across tissues improves, including refinement of methods and gene models, as well as availability of long-read RNA-seq data from human tissues. In addition, improvement of single-cell RNA-seq technologies and generation of data across human tissues will provide insight into cell type-specific exon usage for incorporation into variant interpretation^27^.

The code used to generate pext is available as open source software (https://github.com/macarthur-lab/tx_annotation). In addition, we provide a precomputed file of the transcript expression value for every possible single nucleotide variant in the human genome. This metric has already proven useful in variant curation for drug target identification (*32*) and for filtering variants for identification of human knockouts(*1*). Overall, our metric can be incorporated into variant interpretation in Mendelian disease pipelines, rare variant burden analyses, and the prioritization of variants for recall-by-genotype studies.

## Methods

### Data and code availability

We utilized the gnomAD v.2.1.1 sites Hail 0.2 (https://hail.is) table which is accessible publicly at gs://gnomad-public/release/2.1.1 and at https://gnomad.broadinstitute.org. The GTEx v7 gene and isoform expression data were downloaded from the GTEx portal (gtexportal.org). The GTEx pipeline for isoform quantification is available publicly (https://github.com/broadinstitute/gtex-pipeline/) and briefly involves 2-pass alignment with STAR v2.4.2a(*33*), gene expression quantification with RNA-SeQC v1.1.8(*34*), and isoform quantification with RSEM v1.2.22. The LOEUF constraint file was downloaded from gs://gnomad-resources/lof_paper/. Variants used in all gnomAD analyses in the manuscript passed random forest filtering, and all pLoF variants were annotated as high confidence (HC) by LOFTEE v.1.0, which is described in an accompanying manuscript(*1*). All files used in the analyses in the manuscript are available in gs://gnomad-public/papers/2019-tx-annotation/. Scripts to QC the gnomAD dataset are available at https://github.com/macarthur-lab/gnomad_qc and the scripts to generate files for the analyses are available at https://github.com/macarthur-lab/tx_annotation.

### Curation of pLoF variants in haploinsufficient developmental disease genes

For identification of haploinsufficient developmental delay genes, we selected genes curated by the ClinGen Dosage Sensitivity Working Group(*35*); 58 of the 61 genes had a score of 3 with sufficient evidence for pathogenicity, while two genes (*CHAMP1, CTCF*) had a score of 2 (some evidence) and one gene (*RERE*) was not yet scored. The penetrance of pathogenic variants in each gene was reviewed in the literature, and only genes with >75% reported penetrance were included. These conditions are those too severe to expect to see an individual in gnomAD (likely unable to consent for a study without guardianship). The 61 genes include 50 autosomal genes of high severity and high penetrance and 11 genes on chromosome X where the phenotype is expected to be severe or lethal in males and moderate to severe in females. The resulting gene list is available at gs://gnomad-public/papers/2019-tx-annotation/data/gene_lists/HI_genes_100417.tsv.

We extracted pLoF variants, defined as essential splice acceptor, essential splice donor, stop gained, and frameshift variants, identified in the 61 haploinsufficient disease genes from the gnomAD v2.1.1 exome and genome sites tables, and considered only those pLoF variants that passed random forest filtering in the gnomAD dataset, and were annotated as high confidence (HC) by LOFTEE v1.0. Of 61 genes, 55 had at least one high quality pLoF available in gnomAD. We performed manual curation of 401 pLoF variants using a web-based curation portal to identify any reason a pLoF may have been a variant calling or annotation error, and categorized the likelihood of each variant being a true LoF.

Evidence for classifying an LoF variant as artifactual was categorized into the following groups: mapping error, strand bias, reference error, genotyping error, homopolymer sequence, in-frame multi-nucleotide variant or frame-restoring indel, essential splice site rescue, minority of transcripts, weak exon conservation, last exon, and other annotation error. All possible reasons talso o reject a LoF consequence were flagged, even when a single criterion would categorize the variant as not LoF. Variants were then categorized as LoF, likely LoF, likely not LoF, and not LoF based on criteria outlined in Supplementary Table 2. Supplementary Figure 1A shows the distribution of the LoF verdicts for the 401 pLoF variants.

Technical errors comprised genotyping errors, strand biases, reference errors, and repetitive regions that could be detected by visual inspection of reads in the Integrative Genomics Viewer(*36*) (IGV) and from the UCSC genome browser(*37*). Genotyping errors comprised skewed allele balances (conservative cutoff of ≤35%), low complexity sequences, GC rich regions, homopolymer tracts (≥ 6 base pairs or ≥ 6 trinucleotide repeats) and low quality metrics (genotype quality, or GQ, < 20). Strand bias was flagged when a variant was skewed preferentially on the forward or reverse strand, or when the majority (>90%) of a given strand covered a region; this was often observed around intron/exon boundaries. Strand biases despite balanced coverage of the forward and reverse strands were weighted towards likely not LoF, whereas a strand bias due to skewed strand coverage were weighted alongside other genotyping errors. Reference errors were uncommon, but identified by a small deletion in a given exon, posing as a <5 base pair intron. Most genotyping errors and strand biases in isolation were not deemed critical in deciding whether a variant was likely not LoF or not LoF, with the exception of allele balance ≤25%. Mapping errors were often identified by an enrichment of complex variation surrounding a variant of interest. Furthermore, the UCSC browser was used to highlight mapping discrepancies, such as self-chain alignments, segmental duplications, simple tandem repeats, and microsatellite regions.

In-frame multi-nucleotide variants (MNVs), essential splice site rescue, and frame-restoring insertion-deletions are rescue events that are predicted to restore gene function. MNVs were visualized in IGV and cross checked with codons from the UCSC browser; in frame MNVs that rescued stop codons were scored as not LoF. Essential splice site rescue occurs when an in frame alternative donor or acceptor site is present, which likely has a minimal effect on the transcript. Thirty-six base pairs upstream and downstream of the splice variant were assessed for splice site rescue. Cryptic splice sites within 6 base pairs of the splice variant were considered a complete rescue, rendering the variant not LoF. Rescue sites > 6 base pairs away but within +/- 20 base pairs were weighted with less confidence, scoring as likely not LoF. All potential splice site rescues were validated using Alamut v.2.11 (https://www.interactive-biosoftware.com/alamut-visual/). Frame-restoring indels were identified by scanning approximately +/- 80 base pairs from the annotated indel and counting any insertions/deletions to assess if the frame would be restored.

Transcript errors encompass issues surrounding alternative transcripts, variants within a terminal coding exon, poorly conserved exons, and re-initiation events. Coding variants that occupied the minority (<50%) of NCBI coding RefSeq transcripts for a given gene were considered not LoF. These variants often affected poorly conserved exons, as determined by PhyloP(*38*), PhyloCSF(*19*), and visualization in the UCSC browser(*37*). The only exception to the minority of transcript criteria were cases where the exon was well conserved, which relegated the categorization to likely not LoF. Variants within the last coding exon, or within 50 base pairs of the penultimate coding exon were also considered not LoF, unless 25% < x <50% of the coding sequence was affected, in which case the variant was deemed likely not LoF. If >50% of the coding sequence was disrupted by a variant in the last exon, this was deemed likely LoF. Other transcript errors included: re-initiation errors; upstream stop codons of a given LoF variant; variants that fell on exactly 50% of coding RefSeq transcripts; and/or partial exon conservation. Re-initiation events were flagged when a methionine downstream of the variant in the first coding exon was predicted to restart transcription, and were predicted to be likely not LoF. Variants occurring after a stop codon in the last coding exon were considered not LoF, particularly across the region of the exon or transcript in question. Error categories were grouped for Figure 1 as follows: Minority of transcripts and weak exon conservation were grouped as transcript errors, genotyping errors and homopolymers as sequencing errors, essential splice rescue and MNV grouped as rescue and strand bias was included in other annotation errors.

The criteria above were strictly adhered throughout and manual curation was performed by two independent reviewers to ensure maximum consistency and minimize human error. Any discordance in curation was re-curated by both curators together and resolved. Full results of manual curation are available in Supplementary Table 3.

### Calculation of transcript-expression aware annotation

We first imported the GTEx v7 isoform quantifications into Hail and calculated the median expression of every transcript per tissue. This precomputed summary isoform expression matrix is available for GTEx v7 in gs://gnomad-public/papers/2019-tx-annotation/data/GRCH37_hg19/. We also import and annotate a variant file with the Variant Effect Predictor (VEP) version 85(*39*) against Gencode v19(*20*), implemented in Hail with the LOFTEE v1.0 plugin.

We use the transcript consequences VEP field to calculate the sum of isoform expression for variant annotations, i.e. the annotation-level expression across transcripts (ext). For variants that have multiple consequences for one transcript (for example, a SNV that is both a missense and a splice region variant on one transcript) we use the worst consequence, ordered by VEP (in this example, missense takes precedence over splice region). We filter the consequences to those only occurring on protein coding transcripts. Full ordering of the VEP consequences is available at: useast.ensembl.org/info/genome/variation/prediction/predicted_data.html

We then sum the expression of every transcript per variant, for every combination of consequence, LOFTEE filter, and LOFTEE flag for every tissue (Supplementary Figure 3A). For example, if a SNV is synonymous on ENST1, a LOFTEE HC stop-gained on ENST3 and ENST4, and LOFTEE low-confidence (LC) stop gained variant on ENST 5 and ENST6, the ext values will be synonymous: ENST1, stop-gained HC: ENST 3 + ENST4, and stop-gained LC: ENST5 + ENST6 per tissue. This can be computed with the tx_annotate() function by setting the tx_annotation_type to “expression”. We foresee the non-normalized ext values to be useful when only considering one tissue of interest.

To allow for taking average expression values across tissues of interest, we normalize the expression value for a given value to the total expression of the gene on which the variant is found. This is carried out by dividing the ext value with the sum of the expression of all transcripts per tissue in transcripts-per-million (TPM) v1.1.8(*34*) (Supplementary Figure 3B). The resulting pext (proportion expression across transcript) value can be interpreted as the proportion of the total transcriptional output from a gene that would be affected by the given variant annotation in question. If the gene expression value (and thus the denominator) in a given tissue is 0, the pext value will not be available for that tissue.

We note that for a minority of genes, when RSEM assigns higher relative expression to non-coding transcripts, the sum of the value of coding transcripts can be much smaller than the gene expression value for the transcript, resulting in low pext scores for all coding variants in the gene, and thus resulting in possible filtering of all variants for a given gene. In many cases this appears to be the result of spurious non-coding transcripts with a high degree of exon overlap with true coding transcripts. To prevent this artifact from affecting our analyses, we first calculated the maximum pext score for all variants across all protein coding genes, and removed any gene where the maximum pext score was below 0.2. This resulted in the filtering of 668 genes, representing 3.3% of all genes analyzed. We note that there is no overlap with the 668 genes and the haploinsufficient gene list, 97 of the filtered genes are present in OMIM (representing 1.5% of the OMIM gene list) and 42 filtered genes are considered constrained (representing 1.4% of LOEUF < 0.35, or constrained, genes) thus having low impact on variant interpretation in the context of disease associations.

When taking averages across tissues, such unavailable pext values are not considered (ie. when taking the mean across tissues, we remove NAs). This value can be computed with the tx_annotate() function by setting the tx_annotation_type to “proportion”. For the analyses in this manuscript, we remove reproduction-associated GTEx tissues (endocervix, ectocervix, fallopian tube, prostate, uterus, ovary, testes, vagina), cell lines (transformed fibroblasts, transformed lymphocytes) and any tissue with less than one hundred samples (bladder, brain Cervicalc-1 spinal cord, brain substantia nigra, kidney cortex, minor salivary gland) resulting in the use of 38 GTEx tissues.

The full transcript-expression aware annotation pipeline, implemented in Hail 0.2, is fully available at https://github.com/macarthur-lab/tx_annotation with commands laid out for analyses in the manuscript. Passing a Hail table through the tx_annotate() function returns the same table with a new field entitled “tx_annotation” which provides either the ext or pext value per variant-annotation pair, depending on parameter choice. We provide a helper function to extract the worst consequence and the associated expression values for these annotations. All analyses in the manuscript are based on the worst consequence of variant, ordered by VEP(*39*).

### Functional validation of transcript-expression aware annotation

Conservation analysis was performed using phyloCSF scores using the same file utilized for the LOFTEE plugin, available publicly in gs://gnomad-public/papers/2019-tx-annotation/data/other_data/phylocsf_data.tsv.bgz. We denoted exons with a phyloCSF max open reading frame (ORF) score > 1000 as highly conserved and those with phyloCSF max ORF score < −100 as lowly conserved (Supplementary Figure 5A) and evaluated their average usage in GTEx.

Using the base-level pext values that are used in the gnomAD browser, we filtered to intervals with high or low conservation, and calculated the average pext value in the interval. To evaluate regions with low conservation but high expression, we identified genes harboring unconserved regions with the pext value > 0.9 for pathway enrichment analysis and used the web browser for FUMA GENE2FUNC feature(*40*), which incorporates Reactome(*41*), KEGG(*42*), Gene Ontology(*43*) (GO) as well as other ontologies. Default parameters were used for FUMA, with all protein coding genes as the background list. Results from FUMA pathway analysis are available in Supplementary Figure 12, and full results are available in Supplementary Table 5.

Analysis of pext values for LOFTEE flags and the MAPS calculation were performed utilizing the gnomAD v2.1.1 exome dataset. Calculation of MAPS scores was previously described in Lek et al. 2016(*21*) and is implemented as a Hail module, as described in Karczewski et al. 2019(*1*). MAPS is a relative metric, and cannot be compared across datasets, but is a useful summary metric for the frequency spectrum, indicating deleteriousness as inferred from rarity of variation (high values of MAPS correspond to lower frequency, suggesting the action of negative selection at more deleterious sites). The MAPS scores were calculated on the gnomAD v.2.1.1 dataset partitioning upon the LOEUF score and expression bin. The script for generating MAPs scores is available in the tx-annotation Github repository under /analyses/maps/maps_submit_per_class.py

### Manual evaluation of unexpressed regions in haploinsufficient developmental delay genes using the GENCODE workflow

As an orthogonal evaluation of regions flagged as unexpressed with the pext metric, we identified any region in 61 haploinsufficient disease genes with a mean pext value < 0.1 in all GTEx tissues and in GTEx brain samples, due to the relevance of brain tissues for these disorders, regardless of mutational burden in gnomAD. The resulting list of 128 regions was evaluated by the HAVANA manual annotation group of the GENCODE project(*20*).

The manual evaluation first established whether the transcript model corresponding to the region in question was correct in terms of structure, comparing exon / intron combinations, and the accuracy of splice sites against the RNA evidence supporting the model. Second, the functional biotype of each model was reassessed; in particular, whether the decision to annotate the model as protein-coding in GENCODE v19 was appropriate. Note that GENCODE models that incorporate alternative exons or exon combinations in comparison to the ‘canonical’ isoform are likely to be annotated as coding if they contain a prospective CDS that is considered biologically plausible, based on a mechanistic view of translation. These re-annotations are summarized in Supplementary Table 4.

We binned cases into three main categories, according to confidence in both the accuracy and potential functional relevance of the overlapping models: (1) ‘error’, where the model was seen to have an incorrect transcript structure and/or a CDS that conflicted with updated GENCODE annotation criteria (these annotations had been or will be changed in future GENCODE releases based on this evaluation); (2) ‘putative’, where the model structure and CDS satisfied our current annotation criteria, although we judged the potential of the transcript represented to encode a protein with a functional role in cellular physiology to be nonetheless speculative (these have been maintained as putative protein-coding transcripts in GENCODE); (3) ‘validated’, where we believe it is highly probable that the model represents a true protein-coding isoform. High confidence in the validity of the CDS was based on comparative annotation, i.e. the observation of CDS conservation and also the existence of equivalent transcript models in other species. GENCODE also annotates transcript models as ‘nonsense-mediated decay (NMD) and ‘non-stop decay’ (NSD), where a translation is found that is predicted to direct the RNA molecule into cellular degradation programs. While it has been established that such ‘non-productive’ transcription events can play a role in gene regulation and thus disease, the interpretation of variants within NMD and NSD CDS remains challenging(*44*). These models were therefore classed in a separate category.

### Gene list comparisons

To evaluate the filtering power of the pext metric for Mendelian variants, we evaluated the number of variants that would be filtered with an average GTEx pext cutoff of 0.1 (low expression) in the ClinVar and gnomAD datasets. We downloaded the ClinVar VCF from the ClinVar FTP (version dated 10/28/2018), imported it into Hail, annotated it with VEP v85 against Gencode v19, and added pext annotations with the tx_annotate() function. All evaluated variants were annotated as HC by LOFTEE v1.0, and ClinVar variants were filtered to those marked as pathogenic, with no conflicts, and reviewed with at least one star status.

For variants in 61 haploinsufficient genes, we identified any variant identified in at least one individual with any zygosity in both datasets. For variants identified in autosomal recessive disease genes, we used a list of 1,183 OMIM disease genes deemed to follow a recessive inheritance pattern by Blekhman et al.(*45*) and Berg et al.(*46*) (available as https://github.com/macarthur-lab/gene_lists/blob/master/lists/all_ar.tsv). We compared the pext value for all pLoF variants identified in ClinVar versus any variant in a homozygous state in at least one individual in the gnomAD exome or genome datasets. Finally, we used a LOEUF cutoff of 0.35 to denote constrained genes, and compared any synonymous or pLoF variant in these genes in the gnomAD exome or genome datasets.

### *De novo* and rare variant analysis

De novo variants were collated from previously published studies. We collected *de novo* variants identified in 5,305 probands from trio studies of intellectual disability/developmental disorders (Hamdam et al(*27*): n = 41, de Ligt et al(*28*): N = 100, Rauch et al(*29*): N = 51, DDD(*24*): n = 4,293, Lelieveld et al(*26*): n = 820), 1,073 probands with congenital heart disease with co-morbid developmental delay (Sifrim et al(*42*): n = 512, Chih Jin et al(*47*) : 561), 6,430 ASD probands, and 2,179 unaffected controls from the Autism Sequencing Consortium(*25*). We also utilized a previously published dataset of variants in 8,437 cases with ASD and/or attention-deficit/hyperactivity disorder and 5,214 controls from the Danish Neonatal Screening Biobank (*48*). In this analysis, we analyzed pLoF variants identified in highly constrained genes (first LOEUF decile) with a combined total allele count of ≤10 in cases and controls.

We annotated both de novo and rare variants with VEP v85 against Gencode v19 and added pext annotations with the tx_annotate() function. We then calculated the average pext metric across 11 GTEx brain samples and binned them as low (pext < 0.1), medium (0.1 ≤ pext ≤ 0.9) or high (pext > 0.9) expression. We then calculated the number of pLoF, missense, and synonymous variants per pext expression bin. To obtain case-control rate ratios and the 95% confidence intervals for *de novo* variant analyses, we used a two-sided Poisson exact test on counts(*49*). To obtain the odds ratio for the rare variant analysis in ASD/ADHD, we used the Fisher’s exact test for count data.

### Isoform quantifications via salmon

To evaluate whether use of a different isoform quantification tool would affect results, we compared results of *TCF4* base-level expression (shown in Figure 2B), MAPS (Figure 3C) and comparison of the number of variants filtered in haploinsufficient developmental disease genes in ClinVar vs gnomAD (Figure 4A) using RSEM quantifications used in this study with quantifications using salmon v.0.12(*50*). Due to the intractability of re-quantifying the entire GTEx dataset, we downloaded and requantified 151 GTEx brain – cortex CRAM files from the V7 dataset. We first converted CRAMs to fastq files using Picard 2.18.20 and ran salmon with the “salmon quant –i index -fastq1 – fastq2 –minAssignedFrag1 –validateMappings” command. The index was created with the “salmon index –t transcript.fa –type quasi –k 31” command using the GENCODE v19 protein-coding and lncRNA transcripts FASTA files. The existing GTEx RSEM isoform quantifications were filtered to the same GTEx brain – cortex samples. For the analyses to remain consistent with the remainder of the manuscript, we calculated the maximum Brain - Cortex pext score for all variants across all protein coding genes for both the RSEM and salmon quantifications, and removed any gene where the maximum pext score was below 0.2. This resulted in filtering 325 genes from the salmon quantification of the Brain – Cortex samples and 691 genes from the RSEM quantification, corresponding to 3.4 and 1.6% of quantified genes, respectively. We filtered these genes in both the MAPs and gene list comparison analysis seen in Supplementary Figure 11. The WDL script for the quantification pipeline is available at: gs://gnomad-public/papers/2019-tx-annotation/results/salmon_rsem/salmon.wdl and the commands to obtain results for each individual analysis in the tx-annotation Github repository under /analyses/rsem_salmon/.

### Transcript expression aware annotation with fetal isoform expression dataset

While our analyses were based on transcript expression aware annotation from the GTEx v7 dataset, we provide necessary files for pext annotation with the Human Brain Development Resource (HBDR) fetal brain dataset(*51*) in gs://gnomad-public/papers/2019-tx-annotation/data/HBDR_fetal_RNAseq. HBDR includes 558 samples from varying brain subregions across developmental time points. We downloaded HDBR sample fastq files from European Nucleotide Archive (study accession PRJEB14594) and obtained RSEM isoform quantification on HBDR fastqs using the GTEx v7 quantification pipeline, publicly available at https://github.com/broadinstitute/gtex-pipeline/) which briefly involves 2-pass alignment with STAR v2.4.2a(*33*) and isoform quantification with RSEM v1.2.22. Here, we also removed genes where the average pext across HBDR was below 0.2, resulting in the removal of 712 genes (3.5% of all analyzed genes).The dataset was also used for the analysis of baselevel expression values in *SCN2A* shown in Supplementary Figure 7D.

## Supporting information

upplementary Figure 4 - Baselevel TCF4 expression per GTEx tissue

Supplementary Table 4 - GENCODE curation results of 128 regions flagged as unexpressed by pext

Supplementary Table 3 - Manual curation results of 401 pLoFs in 61 HI developmental disease genes identified in gnomAD

Supplementary Table 5 - FUMA GENE2FUNC analysis results and run information

## Supplementary Materials and Methods

**Fig S1:** Details of manual curation of 401 pLoF variants in 61 HI developmental disease genes

**Fig S2:** Technical artifacts in transcriptome sequencing experiments prevent the use of read pileup at exons as an unbiased proxy for expression

**Fig S3:** Details of calculating transcript-expression annotation

**Fig S4:** Baselevel *TCF4* expression per GTEx tissue

**Fig S5:** Functional validation of pext

**Fig S6:** Results of GENCODE curation of 128 unexpressed regions in HI developmental disease genes

**Fig S7:** Evaluation of pext after accounting for LOFTEE flags in ClinVar and *de novo* analyses

**Fig S8:** Application of pext binning to *de novo* and rare variant analysis in additional datasets

**Fig S9:** Evaluation of using differing cutoffs for average, minimum or maximum pext across tissues

**Fig S10:** Using pext based on fetal RNA-seq isoform expression data for filtering variants in haploinsufficient genes

**Fig S11:** Comparison of key results using Salmon vs RSEM.

**Fig S12:** Results from FUMA GENE2FUNC analysis in unconserved regions with high expression values

**Table S1:** Summary of manual curation flags for 401 pLoFs in 61 HI developmental disease genes identified in gnomAD

**Table S2:** Summary of criteria for LoF verdicts of 401 pLoF in 61 HI developmental disease genes identified in gnomAD

**Table S3:** Manual curation results of 401 pLoFs in 61 HI developmental disease genes identified in gnomAD

**Table S4:** GENCODE curation results of 128 regions flagged as unexpressed by pext

**Table S5:** FUMA GENE2FUNC analysis results and run information

## Acknowledgements

We thank all of the research participants for contributing their data. This work was supported by NIDDK U54 DK105566, NIGMS R01 GM104371, and the Broad Institute. KJK was supported by NIGMS F32 GM115208. A.O.L was supported by NICHD K12 HD052896. The GENCODE project is supported by the National Human Genome Research Institute of the National Institutes of Health under Award Number U41HG007234. The results published here are in part based upon data: 1) generated by The Cancer Genome Atlas managed by the NCI and NHGRI (accession: <u>phs000178.v10.p8</u>). Information about TCGA can be found at http://cancergenome.nih.gov, 2) generated by the Genotype-Tissue Expression Project (GTEx) managed by the NIH Common Fund and NHGRI (accession: <u>phs000424.v7.p2</u>), 3) generated by the Exome Sequencing Project, managed by NHLBI, 4) generated by the Alzheimer’s Disease Sequencing Project (ADSP), managed by the NIA and NHGRI (accession: <u>phs000572.v7.p4</u>). We thank Emma Pierce-Hoffman for prior analysis and thoughts on characterizing loss of function variants in haploinsufficient genes. We thank the iPSYCH/SSI/Broad Institute psychiatric genetics study for the use of exome count data.

We have complied with all relevant ethical regulations. This study was overseen by the Broad Institute’s Office of Research Subject Protection and the Partners Human Research Committee, and was given a determination of Not Human Subjects Research. Informed consent was obtained from all participants.

## Group authors

### Genome Aggregation Database Production Team

Jessica Alföldi^1,2^, Irina M. Armean^3,1,2^, Eric Banks^4^, Louis Bergelson^4^, Kristian Cibulskis^4^, Ryan L Collins^1,5,6^, Kristen M. Connolly^7^, Miguel Covarrubias^4^, Beryl Cummings^1,2,8^, Mark J. Daly^1,2,9^, Stacey Donnelly^1^, Yossi Farjoun^4^, Steven Ferriera^10^, Laurent Francioli^1,2^, Stacey Gabriel^10^, Laura D. Gauthier^4^, Jeff Gentry^4^, Namrata Gupta^10,1^, Thibault Jeandet^4^, Diane Kaplan^4^, Konrad J. Karczewski^1,2^, Kristen M. Laricchia^1,2^, Christopher Llanwarne^4^, Eric V. Minikel^1^, Ruchi Munshi^4^, Benjamin M Neale^1,2^, Sam Novod^4^, Anne H. O’Donnell-Luria^1,11,12^, Nikelle Petrillo^4^, Timothy Poterba^9,2,1^, David Roazen^4^, Valentin Ruano-Rubio^4^, Andrea Saltzman^1^, Kaitlin E. Samocha^13^, Molly Schleicher^1^, Cotton Seed^9,2^, Matthew Solomonson^1,2^, Jose Soto^4^, Grace Tiao^1,2^, Kathleen Tibbetts^4^, Charlotte Tolonen^4^, Christopher Vittal^9,2^, Gordon Wade^4^, Arcturus Wang^9,2,1^, Qingbo Wang^1,2,6^, James S Ware^14,15,1^, Nicholas A Watts^1,2^, Ben Weisburd^4^, Nicola Whiffin^14,15,1^

1. Program in Medical and Population Genetics, Broad Institute of MIT and Harvard, Cambridge, Massachusetts 02142, USA
2. Analytic and Translational Genetics Unit, Massachusetts General Hospital, Boston, Massachusetts 02114, USA
3. European Molecular Biology Laboratory, European Bioinformatics Institute, Wellcome Genome Campus, Hinxton, Cambridge, CB10 1SD, United Kingdom
4. Data Sciences Platform, Broad Institute of MIT and Harvard, Cambridge, Massachusetts 02142, USA
5. Center for Genomic Medicine, Massachusetts General Hospital, Boston, MA 02114, USA
6. Program in Bioinformatics and Integrative Genomics, Harvard Medical School, Boston, MA 02115, USA
7. Genomics Platform, Broad Institute of MIT and Harvard, Cambridge, Massachusetts 02142, USA
8. Program in Biological and Biomedical Sciences, Harvard Medical School, Boston, MA, 02115, USA
9. Stanley Center for Psychiatric Research, Broad Institute of MIT and Harvard, Cambridge, Massachusetts 02142, USA
10. Broad Genomics, Broad Institute of MIT and Harvard, Cambridge, Massachusetts 02142, USA
11. Division of Genetics and Genomics, Boston Children’s Hospital, Boston, Massachusetts 02115, USA
12. Department of Pediatrics, Harvard Medical School, Boston, Massachusetts 02115, USA
13. Wellcome Sanger Institute, Wellcome Genome Campus, Hinxton, Cambridge CB10 1SA, UK
14. National Heart & Lung Institute and MRC London Institute of Medical Sciences, Imperial College London, London UK
15. Cardiovascular Research Centre, Royal Brompton & Harefield Hospitals NHS Trust, London UK

### Genome Aggregation Database Consortium

Carlos A Aguilar Salinas^1^, Tariq Ahmad^2^, Christine M. Albert^3,4^, Diego Ardissino^5^, Gil Atzmon^6,7^, John Barnard^8^, Laurent Beaugerie^9^, Emelia J. Benjamin^10,11,12^, Michael Boehnke^13^, Lori L. Bonnycastle^14^, Erwin P. Bottinger^15^, Donald W Bowden^16,17,18^, Matthew J Bown^19,20^, John C Chambers^21,22,23^, Juliana C. Chan^24^, Daniel Chasman^3,25^, Judy Cho^15^, Mina K. Chung^26^, Bruce Cohen^27,25^, Adolfo Correa^28^, Dana Dabelea^29^, Mark J. Daly^30,31,32^, Dawood Darbar^33^, Ravindranath Duggirala^34^, Josée Dupuis^35,36^, Patrick T. Ellinor^30,37^, Roberto Elosua^38,39,40^, Jeanette Erdmann^41,42,43^, Tõnu Esko^30,44^, Martti Färkkilä^45^, Jose Florez^46^, Andre Franke^47^, Gad Getz^48,49,25^, Benjamin Glaser^50^, Stephen J. Glatt^51^, David Goldstein^52,53^, Clicerio Gonzalez^54^, Leif Groop^55,56^, Christopher Haiman^57^, Craig Hanis^58^, Matthew Harms^59,60^, Mikko Hiltunen^61^, Matti M. Holi^62^, Christina M. Hultman^63,64^, Mikko Kallela^65^, Jaakko Kaprio^56,66^, Sekar Kathiresan^67,68,25^, Bong-Jo Kim^69^, Young Jin Kim^69^, George Kirov^70^, Jaspal Kooner^23,22,71^, Seppo Koskinen^72^, Harlan M. Krumholz^73^, Subra Kugathasan^74^, Soo Heon Kwak^75^, Markku Laakso^76,77^, Terho Lehtimäki^78^, Ruth J.F. Loos^15,79^, Steven A. Lubitz^30,37^, Ronald C.W. Ma^24,80,81^, Daniel G. MacArthur^31,30^, Jaume Marrugat^82,39^, Kari M. Mattila^78^, Steven McCarroll^32,83^, Mark I McCarthy^84,85,86^, Dermot McGovern^87^, Ruth McPherson^88^, James B. Meigs^89,25,90^, Olle Melander^91^, Andres Metspalu^44^, Benjamin M Neale^30,31^, Peter M Nilsson^92^, Michael C O’Donovan^70^, Dost Ongur^27,25^, Lorena Orozco^93^, Michael J Owen^70^, Colin N.A. Palmer^94^, Aarno Palotie^56,32,31^, Kyong Soo Park^75,95^, Carlos Pato^96^, Ann E. Pulver^97^, Nazneen Rahman^98^, Anne M. Remes^99^, John D. Rioux^100,101^, Samuli Ripatti^56,66,102^, Dan M. Roden^103,104^, Danish Saleheen^105,106,107^, Veikko Salomaa^108^, Nilesh J. Samani^19,20^, Jeremiah Scharf^30,32,67^, Heribert Schunkert^109,110^, Moore B. Shoemaker^111^, Pamela Sklar*^112,113,114^, Hilkka Soininen^115^, Harry Sokol^9^, Tim Spector^116^, Patrick F. Sullivan^63,117^, Jaana Suvisaari^108^, E Shyong Tai^118,119,120^, Yik Ying Teo^118,121,122^, Tuomi Tiinamaija^56,123,124^, Ming Tsuang^125,126^, Dan Turner^127^, Teresa Tusie-Luna^128,129^, Erkki Vartiainen^66^, James S Ware^130,131,30^, Hugh Watkins^132^, Rinse K Weersma^133^, Maija Wessman^123,56^, James G. Wilson^134^, Ramnik J. Xavier^135,136^

1. Unidad de Investigacion de Enfermedades Metabolicas. Instituto Nacional de Ciencias Medicas y Nutricion. Mexico City
2. Peninsula College of Medicine and Dentistry, Exeter, UK
3. Division of Preventive Medicine, Brigham and Women’s Hospital, Boston, Massachusetts, USA.
4. Division of Cardiovascular Medicine, Brigham and Women’s Hospital and Harvard Medical School, Boston, Massachusetts, USA.
5. Department of Cardiology, University Hospital, 43100 Parma, Italy
6. Department of Biology, Faculty of Natural Sciences, University of Haifa, Haifa, Israel
7. Departments of Medicine and Genetics, Albert Einstein College of Medicine, Bronx, NY, USA, 10461
8. Department of Quantitative Health Sciences, Lerner Research Institute, Cleveland Clinic, Cleveland, OH 44122, USA
9. Sorbonne Université, APHP, Gastroenterology Department, Saint Antoine Hospital, Paris, France
10. NHLBI and Boston University’s Framingham Heart Study, Framingham, Massachusetts, USA.
11. Department of Medicine, Boston University School of Medicine, Boston, Massachusetts, USA.
12. Department of Epidemiology, Boston University School of Public Health, Boston, Massachusetts, USA.
13. Department of Biostatistics and Center for Statistical Genetics, University of Michigan, Ann Arbor, Michigan 48109
14. National Human Genome Research Institute, National Institutes of Health, Bethesda, MD, USA
15. The Charles Bronfman Institute for Personalized Medicine, Icahn School of Medicine at Mount Sinai, New York, NY
16. Department of Biochemistry, Wake Forest School of Medicine, Winston-Salem, NC, USA
17. Center for Genomics and Personalized Medicine Research, Wake Forest School of Medicine, Winston-Salem, NC, USA
18. Center for Diabetes Research, Wake Forest School of Medicine, Winston-Salem, NC, USA
19. Department of Cardiovascular Sciences, University of Leicester, Leicester, UK
20. NIHR Leicester Biomedical Research Centre, Glenfield Hospital, Leicester, UK
21. Department of Epidemiology and Biostatistics, Imperial College London, London, UK
22. Department of Cardiology, Ealing Hospital NHS Trust, Southall, UK
23. Imperial College Healthcare NHS Trust, Imperial College London, London, UK
24. Department of Medicine and Therapeutics, The Chinese University of Hong Kong, Hong Kong, China.
25. Department of Medicine, Harvard Medical School, Boston, MA
26. Departments of Cardiovascular Medicine, Cellular and Molecular Medicine, Molecular Cardiology, and Quantitative Health Sciences, Cleveland Clinic, Cleveland, Ohio, USA.
27. McLean Hospital, Belmont, MA
28. Department of Medicine, University of Mississippi Medical Center, Jackson, Mississippi, USA
29. Department of Epidemiology, Colorado School of Public Health, Aurora, Colorado, USA.
30. Program in Medical and Population Genetics, Broad Institute of MIT and Harvard, Cambridge, MA, USA
31. Analytic and Translational Genetics Unit, Massachusetts General Hospital, Boston, Massachusetts 02114, USA
32. Stanley Center for Psychiatric Research, Broad Institute of MIT and Harvard, Cambridge, MA, USA
33. Department of Medicine and Pharmacology, University of Illinois at Chicago
34. Department of Genetics, Texas Biomedical Research Institute, San Antonio, TX, USA
35. Department of Biostatistics, Boston University School of Public Health, Boston, MA 02118, USA
36. National Heart, Lung, and Blood Institute’s Framingham Heart Study, Framingham, MA 01702, USA
37. Cardiac Arrhythmia Service and Cardiovascular Research Center, Massachusetts General Hospital, Boston, MA
38. Cardiovascular Epidemiology and Genetics, Hospital del Mar Medical Research Institute (IMIM). Barcelona, Catalonia, Spain
39. CIBER CV, Barcelona, Catalonia, Spain
40. Departament of Medicine, Medical School, University of Vic-Central University of Catalonia. Vic, Catalonia, Spain
41. Institute for Cardiogenetics, University of Lübeck, Lübeck, Germany
42. 1. DZHK (German Research Centre for Cardiovascular Research), partner site Hamburg/Lübeck/Kiel, 23562 Lübeck, Germany
43. University Heart Center Lübeck, 23562 Lübeck, Germany
44. Estonian Genome Center, Institute of Genomics, University of Tartu, Tartu, Estonia
45. Helsinki University and Helsinki University Hospital, Clinic of Gastroenterology, Helsinki, Finland.
46. Diabetes Unit and Center for Genomic Medicine, Massachusetts General Hospital; Programs in Metabolism and Medical & Population Genetics, Broad Institute; Department of Medicine, Harvard Medical School
47. Institute of Clinical Molecular Biology (IKMB), Christian-Albrechts-University of Kiel, Kiel, Germany
48. Bioinformatics Program, MGH Cancer Center and Department of Pathology
49. Cancer Genome Computational Analysis, Broad Institute.
50. Endocrinology and Metabolism Department, Hadassah-Hebrew University Medical Center, Jerusalem, Israel
51. Department of Psychiatry and Behavioral Sciences; SUNY Upstate Medical University
52. Institute for Genomic Medicine, Columbia University Medical Center, Hammer Health Sciences, 1408, 701 West 168th Street, New York, New York 10032, USA.
53. Department of Genetics & Development, Columbia University Medical Center, Hammer Health Sciences, 1602, 701 West 168th Street, New York, New York 10032, USA.
54. Centro de Investigacion en Salud Poblacional. Instituto Nacional de Salud Publica MEXICO
55. Lund University, Sweden
56. Institute for Molecular Medicine Finland (FIMM), HiLIFE, University of Helsinki, Helsinki, Finland
57. Lund University Diabetes Centre
58. Human Genetics Center, University of Texas Health Science Center at Houston, Houston, TX 77030
59. Department of Neurology, Columbia University
60. Institute of Genomic Medicine, Columbia University
61. Institute of Biomedicine, University of Eastern Finland, Kuopio, Finland
62. Department of Psychiatry, PL 320, Helsinki University Central Hospital, Lapinlahdentie, 00 180 Helsinki, Finland
63. Department of Medical Epidemiology and Biostatistics, Karolinska Institutet, Stockholm, Sweden
64. Icahn School of Medicine at Mount Sinai, New York, NY, USA
65. Department of Neurology, Helsinki University Central Hospital, Helsinki, Finland.
66. Department of Public Health, Faculty of Medicine, University of Helsinki, Finland
67. Center for Genomic Medicine, Massachusetts General Hospital, Boston, Massachusetts 02114, USA
68. Cardiovascular Disease Initiative and Program in Medical and Population Genetics, Broad Institute of MIT and Harvard, Cambridge, Massachusetts 02142, USA
69. Center for Genome Science, Korea National Institute of Health, Chungcheongbuk-do, Republic of Korea.
70. MRC Centre for Neuropsychiatric Genetics & Genomics, Cardiff University School of Medicine, Hadyn Ellis Building, Maindy Road, Cardiff CF24 4HQ
71. National Heart and Lung Institute, Cardiovascular Sciences, Hammersmith Campus, Imperial College London, London, UK.
72. Department of Health, THL-National Institute for Health and Welfare, 00271 Helsinki, Finland.
73. Section of Cardiovascular Medicine, Department of Internal Medicine, Yale School of Medicine, New Haven, Connecticut3Center for Outcomes Research and Evaluation, Yale-New Haven Hospital, New Haven, Connecticut.
74. Division of Pediatric Gastroenterology, Emory University School of Medicine, Atlanta, Georgia, USA.
75. Department of Internal Medicine, Seoul National University Hospital, Seoul, Republic of Korea
76. The University of Eastern Finland, Institute of Clinical Medicine, Kuopio, Finland
77. Kuopio University Hospital, Kuopio, Finland
78. Department of Clinical Chemistry, Fimlab Laboratories and Finnish Cardiovascular Research Center-Tampere, Faculty of Medicine and Health Technology, Tampere University, Finland
79. The Mindich Child Health and Development Institute, Icahn School of Medicine at Mount Sinai, New York, NY
80. Li Ka Shing Institute of Health Sciences, The Chinese University of Hong Kong, Hong Kong, China.
81. Hong Kong Institute of Diabetes and Obesity, The Chinese University of Hong Kong, Hong Kong, China.
82. Cardiovascular Research REGICOR Group, Hospital del Mar Medical Research Institute (IMIM). Barcelona, Catalonia.
83. Department of Genetics, Harvard Medical School, Boston, MA, USA
84. Oxford Centre for Diabetes, Endocrinology and Metabolism, University of Oxford, Churchill Hospital, Old Road, Headington, Oxford, OX3 7LJ UK
85. Wellcome Centre for Human Genetics, University of Oxford, Roosevelt Drive, Oxford OX3 7BN, UK
86. Oxford NIHR Biomedical Research Centre, Oxford University Hospitals NHS Foundation Trust, John Radcliffe Hospital, Oxford OX3 9DU, UK
87. F Widjaja Foundation Inflammatory Bowel and Immunobiology Research Institute, Cedars-Sinai Medical Center, Los Angeles, CA, USA.
88. Atherogenomics Laboratory, University of Ottawa Heart Institute, Ottawa, Canada
89. Division of General Internal Medicine, Massachusetts General Hospital, Boston, MA, 02114
90. Program in Population and Medical Genetics, Broad Institute, Cambridge, MA
91. Department of Clinical Sciences, University Hospital Malmo Clinical Research Center, Lund University, Malmo, Sweden.
92. Lund University, Dept. Clinical Sciences, Skane University Hospital, Malmo, Sweden
93. Instituto Nacional de Medicina Genómica (INMEGEN), Mexico City, 14610, Mexico
94. Medical Research Institute, Ninewells Hospital and Medical School, University of Dundee, Dundee, UK.
95. Department of Molecular Medicine and Biopharmaceutical Sciences, Graduate School of Convergence Science and Technology, Seoul National University, Seoul, Republic of Korea
96. Department of Psychiatry, Keck School of Medicine at the University of Southern California, Los Angeles, California, USA.
97. Department of Psychiatry and Behavioral Sciences, Johns Hopkins University School of Medicine, Baltimore, Maryland, USA
98. Division of Genetics and Epidemiology, Institute of Cancer Research, London SM2 5NG
99. Medical Research Center, Oulu University Hospital, Oulu, Finland and Research Unit of Clinical Neuroscience, Neurology, University of Oulu, Oulu, Finland.
100. Research Center, Montreal Heart Institute, Montreal, Quebec, Canada, H1T 1C8
101. Department of Medicine, Faculty of Medicine, Université de Montréal, Québec, Canada
102. Broad Institute of MIT and Harvard, Cambridge MA, USA
103. Department of Biomedical Informatics, Vanderbilt University Medical Center, Nashville, Tennessee, USA.
104. Department of Medicine, Vanderbilt University Medical Center, Nashville, Tennessee, USA.
105. Department of Biostatistics and Epidemiology, Perelman School of Medicine at the University of Pennsylvania, Philadelphia, PA, USA
106. Department of Medicine, Perelman School of Medicine at the University of Pennsylvania, Philadelphia, PA, USA
107. Center for Non-Communicable Diseases, Karachi, Pakistan
108. National Institute for Health and Welfare, Helsinki, Finland
109. Deutsches Herzzentrum München, Germany
110. Technische Universität München
111. Division of Cardiovascular Medicine, Nashville VA Medical Center and Vanderbilt University, School of Medicine, Nashville, TN 37232-8802, USA.
112. Department of Psychiatry, Icahn School of Medicine at Mount Sinai, New York, NY, USA
113. Department of Genetics and Genomic Sciences, Icahn School of Medicine at Mount Sinai, New York, NY, USA
114. Institute for Genomics and Multiscale Biology, Icahn School of Medicine at Mount Sinai, New York, NY, USA
115. Institute of Clinical Medicine, neurology, University of Eastern Finad, Kuopio, Finland
116. Department of Twin Research and Genetic Epidemiology, King’s College London, London UK
117. Departments of Genetics and Psychiatry, University of North Carolina, Chapel Hill, NC, USA
118. Saw Swee Hock School of Public Health, National University of Singapore, National University Health System, Singapore
119. Department of Medicine, Yong Loo Lin School of Medicine, National University of Singapore, Singapore
120. Duke-NUS Graduate Medical School, Singapore
121. Life Sciences Institute, National University of Singapore, Singapore.
122. Department of Statistics and Applied Probability, National University of Singapore, Singapore.
123. Folkhälsan Institute of Genetics, Folkhälsan Research Center, Helsinki, Finland
124. HUCH Abdominal Center, Helsinki University Hospital, Helsinki, Finland
125. Center for Behavioral Genomics, Department of Psychiatry, University of California, San Diego
126. Institute of Genomic Medicine, University of California, San Diego
127. Juliet Keidan Institute of Pediatric Gastroenterology, Shaare Zedek Medical Center, The Hebrew University of Jerusalem, Israel
128. Instituto de Investigaciones Biomédicas UNAM Mexico City
129. Instituto Nacional de Ciencias Médicas y Nutrición Salvador Zubirán Mexico City
130. National Heart & Lung Institute & MRC London Institute of Medical Sciences, Imperial College London, London UK
131. Cardiovascular Research Centre, Royal Brompton & Harefield Hospitals NHS Trust, London UK
132. Radcliffe Department of Medicine, University of Oxford, Oxford UK
133. Department of Gastroenterology and Hepatology, University of Groningen and University Medical Center Groningen, Groningen, the Netherlands
134. Department of Physiology and Biophysics, University of Mississippi Medical Center, Jackson, MS 39216, USA
135. Program in Infectious Disease and Microbiome, Broad Institute of MIT and Harvard, Cambridge, MA, USA
136. Center for Computational and Integrative Biology, Massachusetts General Hospital

