## Supplementary Figures for "Transcript expression-aware annotation improves rare variant discovery and interpretation"

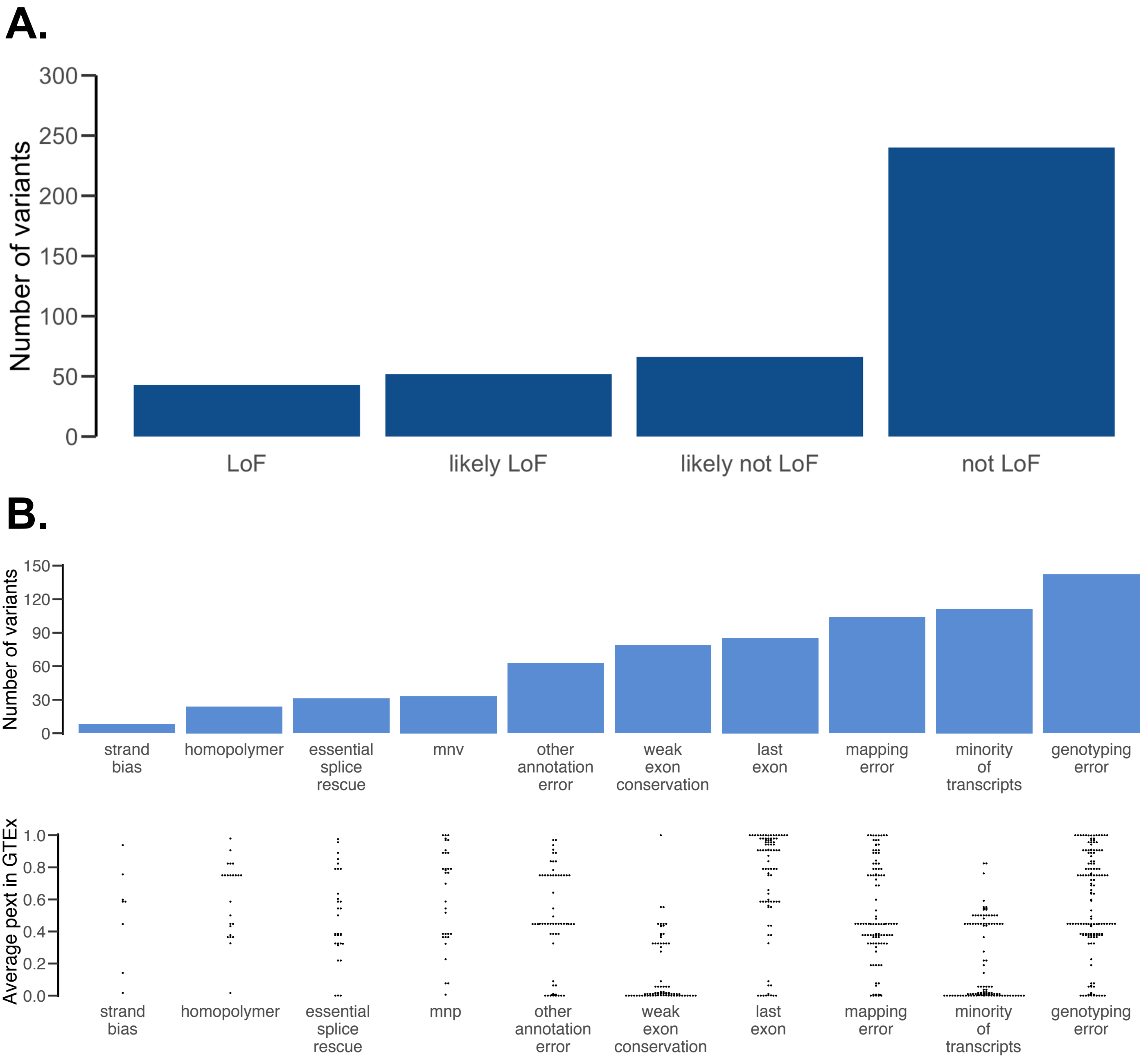


**Supplementary Figure 1: Details of manual curation of 401 pLoF variants in 61 haploinsufficient developmental disease genes A.** Distribution of curation verdicts for the 401 pLoF variants. We categorized 240 variants (76%) as not being LoF, 66 as likely not LoF, 52 as likely LoF and 43 as LoF **B.** Full distribution of the flags refuting true LoF status for 306 not LoF and likely not LoF variants (top) and their corresponding pext score in GTEx (bottom). A variant with multiple flags is assigned to each flag as in Figure 1 (ie. double counted). Minority of transcripts and weak exon conservation were grouped as transcript errors, genotyping errors and homopolymers grouped as sequencing errors, essential splice rescue and MNVs grouped as rescue and strand bias was included in other annotation errors. While the pext values are randomly distributed for other error modes, they are enriched for lower values in transcript errors. Criteria for curation for each flag and verdict, and full curation results are available in Supplementary Tables 1, 2, and 3, respectively.

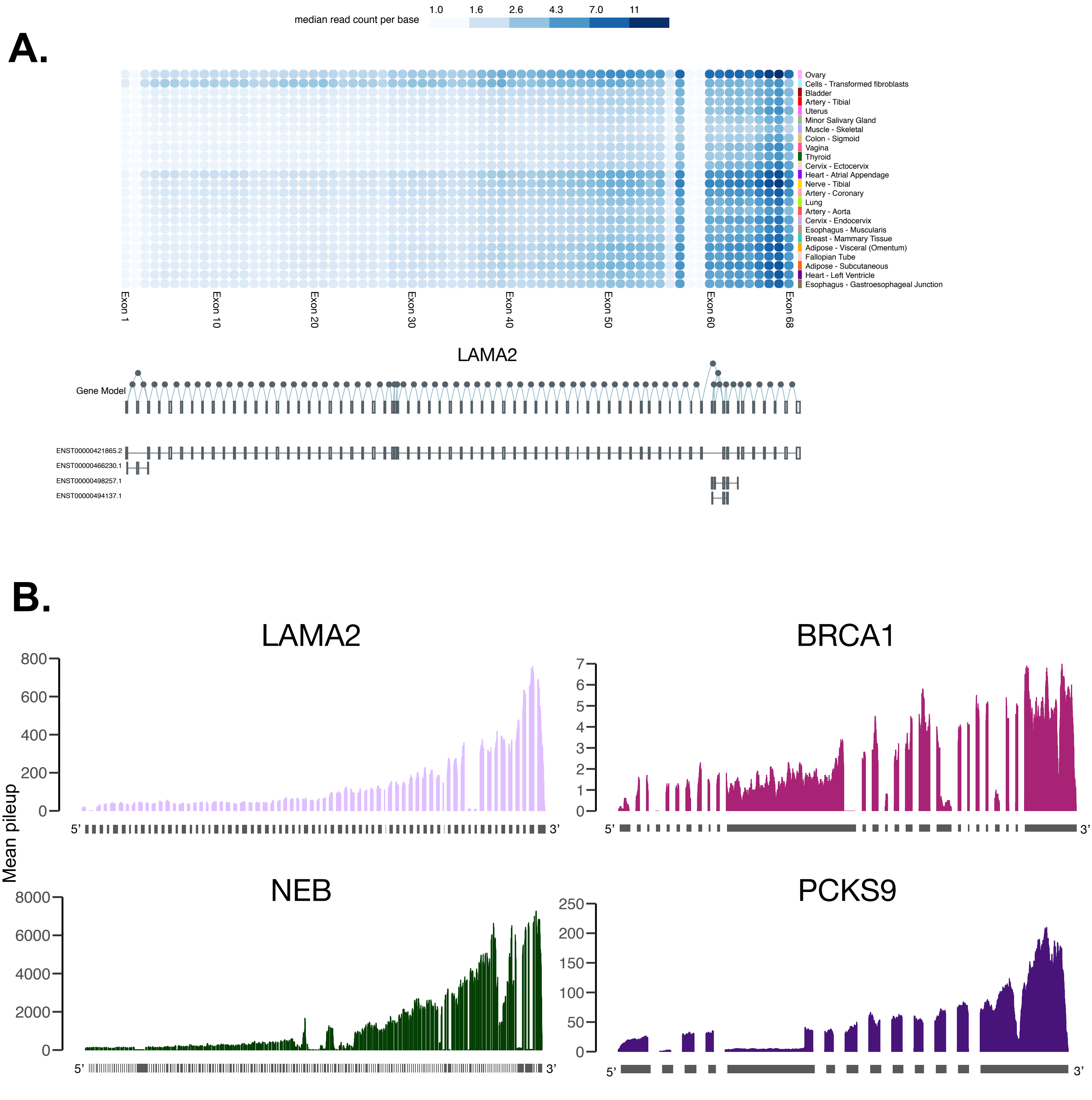


**Supplementary Figure 2: Technical artifacts in transcriptome sequencing experiments prevent the use of read pileup at exons as an unbiased proxy for expression A.** Example of exon expression information on the GTEx web browser (gtexportal.org) for *LAMA2,* which has 3 annotated transcripts. Blue-gray gradient represents median read count per base. While exons 5 and 55 are annotated on a single transcript, the mean read count for the exons are 0.8 and 3.25 in GTEx ovary, respectively, reflecting the confounding effect of 3' bias **B.** Examples of 3' bias in genes of varying lengths and expression levels shows 3' bias is pervasive. Base-level coverage of uniquely mapped reads were calculated in 10 random GTEx samples per tissue using samtools depth in tissues where the genes are highly expressed. Plots show (1) *LAMA2* in tibial nerve (2) *BRCA1* in mammary tissue (breast) (3) *NEB* in skeletal muscle and (4) *PCSK9* in liver, all of which display 3' bias.

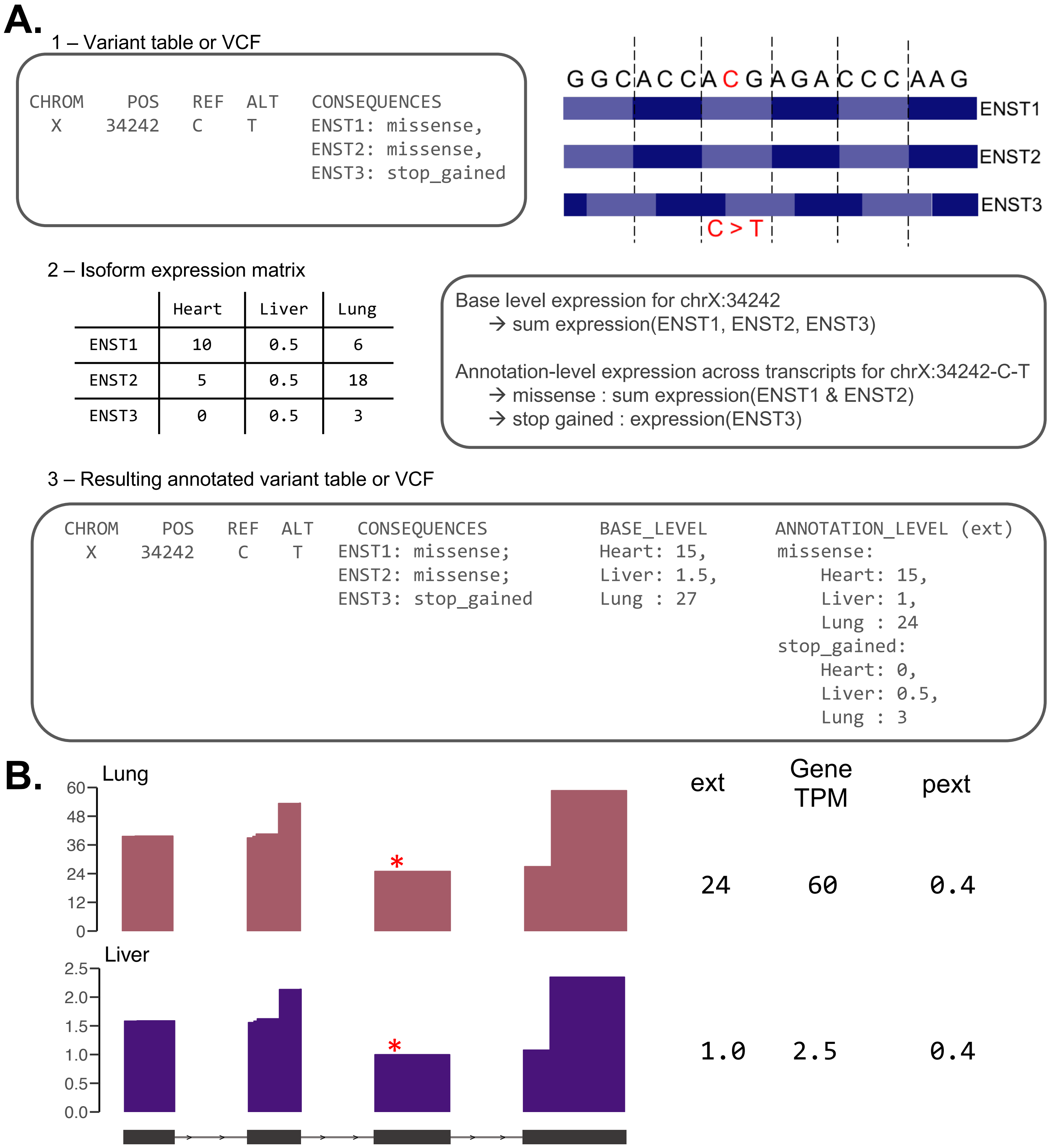


**Supplementary Figure 3: Details of calculating transcript-expression annotation. A.** A SNV can have different consequences across transcripts. For example, an SNV on a region with three annotated transcripts can have a missense effect on two transcripts and a nonsense effect on one transcript. The base-level expression, mainly to be utilized for quick visualization of variant expression in genes, is calculated as the sum of the three transcripts. The annotation level expression across transcript (ext) metric defines the expression of a variant as the sum of the expression of transcripts on which an annotation exists. In this example, the expression value for the missense variant will be the sum of the expression on transcripts where the variant is a missense (ENST1 and ENST2) and the value for the nonsense will the the sum of the expression of transcripts where the variant is a nonsense (ENST3). **B.** To account for gene expression differences between tissues, we normalize the ext value by the sum of expression of all transcript (representing gene expression) in the tissue to calculate the proportion expressed across transcript (pext) value used in the manuscript. This allows for combining pext values across tissues to for example, get the mean pext value across GTEx.

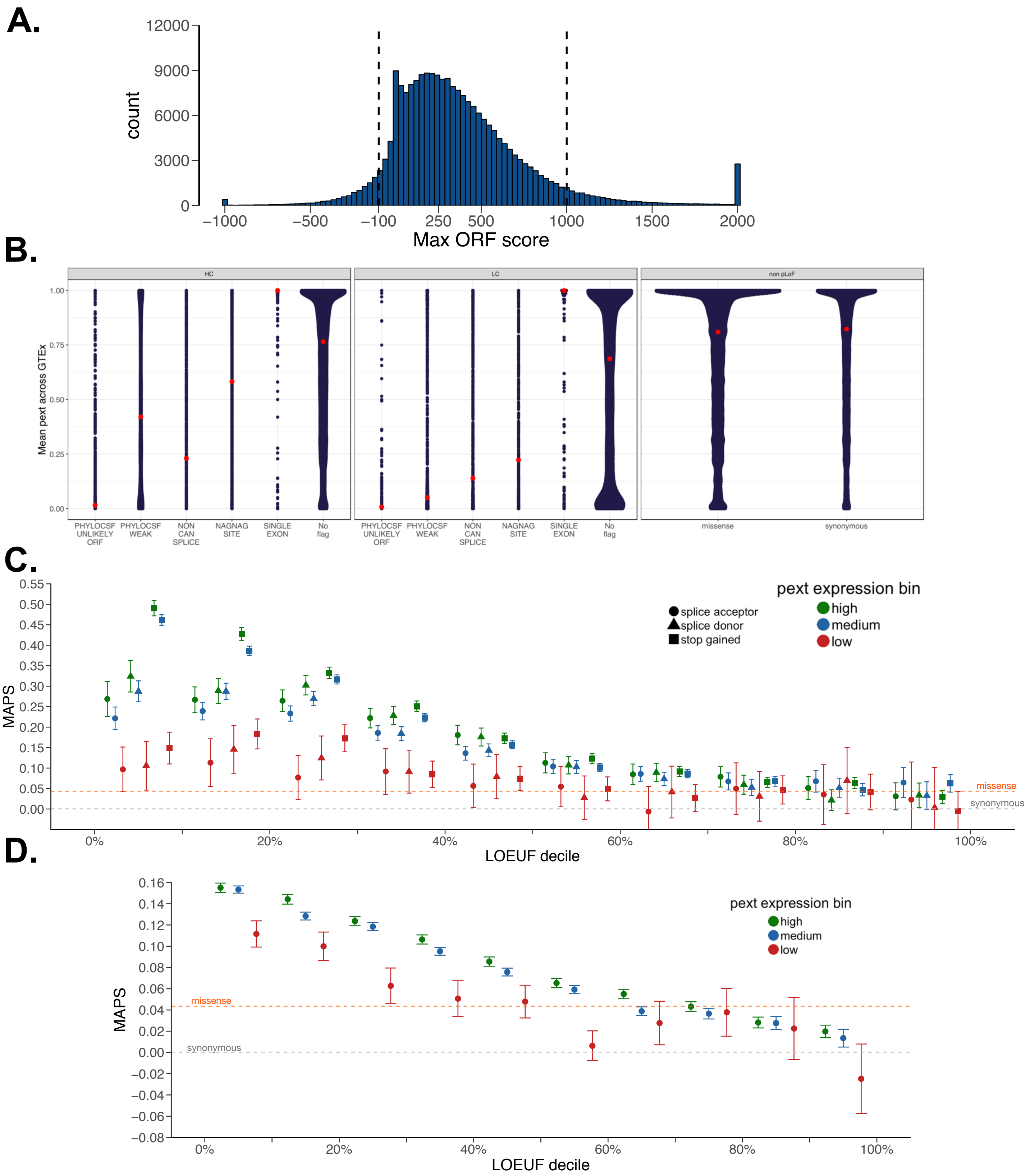


**Supplementary Figure 5: Functional validation of pext A.** Distribution of max open reading frame (ORF) scores from phyloCSF across the genome. We denoted exons with a maximum phyloCSF ORF score > 1000 as highly conserved and those with maximum phyloCSF ORF score < -100 as unconserved. Max ORF scores were capped at -1000 and 2000 for plotting. **B.** Sina plots of pext distribution in all gnomAD exome variants, partitioned on LOFTEE flags and filters (filters denoted as gray bars above plots). Red dots denote median average pext value per category **C.** MAPS score for pLoF variants broken down by specific pLoF consequence shows consistent differences in MAPS for each pLoF category between high, medium and low pext expression bins. **D.** MAPS score of only missense variants denoted probably damaging by Polyphen-2 shows consistent gradient based on high (>0.9) , medium (0.1≤ x ≤ 0.9 ) and low (<0.1) average GTEx pext expression bins showing that pext scores offer supplemental information to existing variant prioritization tools.

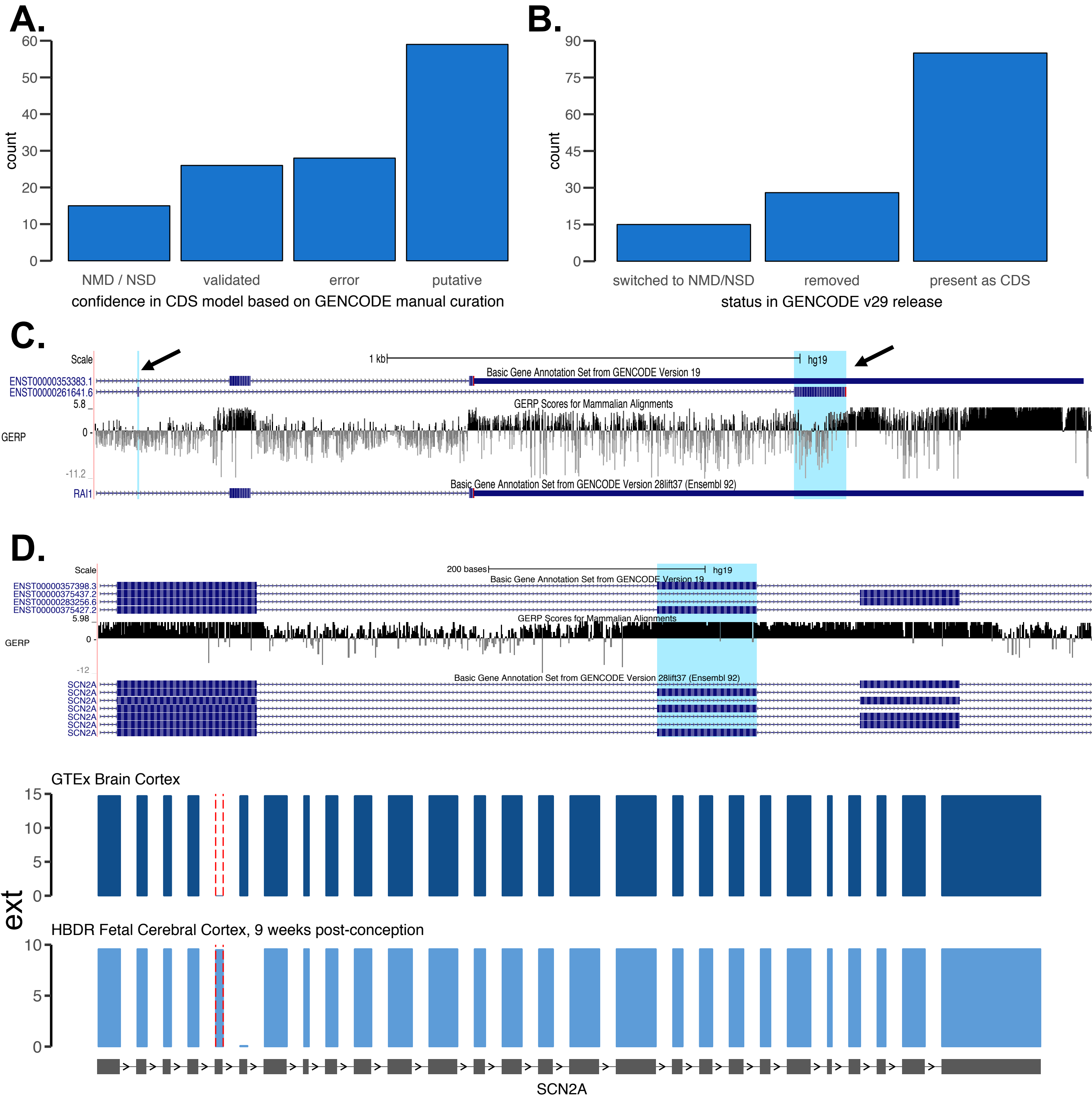


**Supplementary Figure 6: Results of GENCODE of 128 unexpressed regions in haploinsufficient disease genes A.** Summary of confidence in the CDS models tagged as unexpressed in GTEx based on expert manual evaluation. The major curation mode was putative annotation, where regions meet minimal annotation criteria but the coding potential of the region remains speculative. This was followed by regions that were marked as errors and non-coding regions, and have since been removed or are marked for removal based on this analysis. **B.** Summary of current annotation status of the regions in GENCODE v29. While some regions have been removed, or have been switched to a noncoding biotype, a majority remain in subsequent annotation sets **C.** An example of two erroneous gencode v19 CDS' in *RAI1* (chr17:17712481-17712483 and chr17:17714069-17714194, highlighted in blue) flagged by pext as unexpressed. The regions exhibit poor conservation and represents an incorrectly computationally predicted microexon and it's downsteam CDS, likely due to a poor quality cDNA alignment **D.** An example of a likely-coding CDS in *SCN2A* (chr2:166165675-166165766) which is well-conserved (gene model on top). While the region is unexpressed in GTEx, it exhibits considerable expression in fetal tissues from the Human Brain Developmental Resource (shown on bottom), highlighting the importance of incorporating multiple isoform datasets for accurate interpretation.

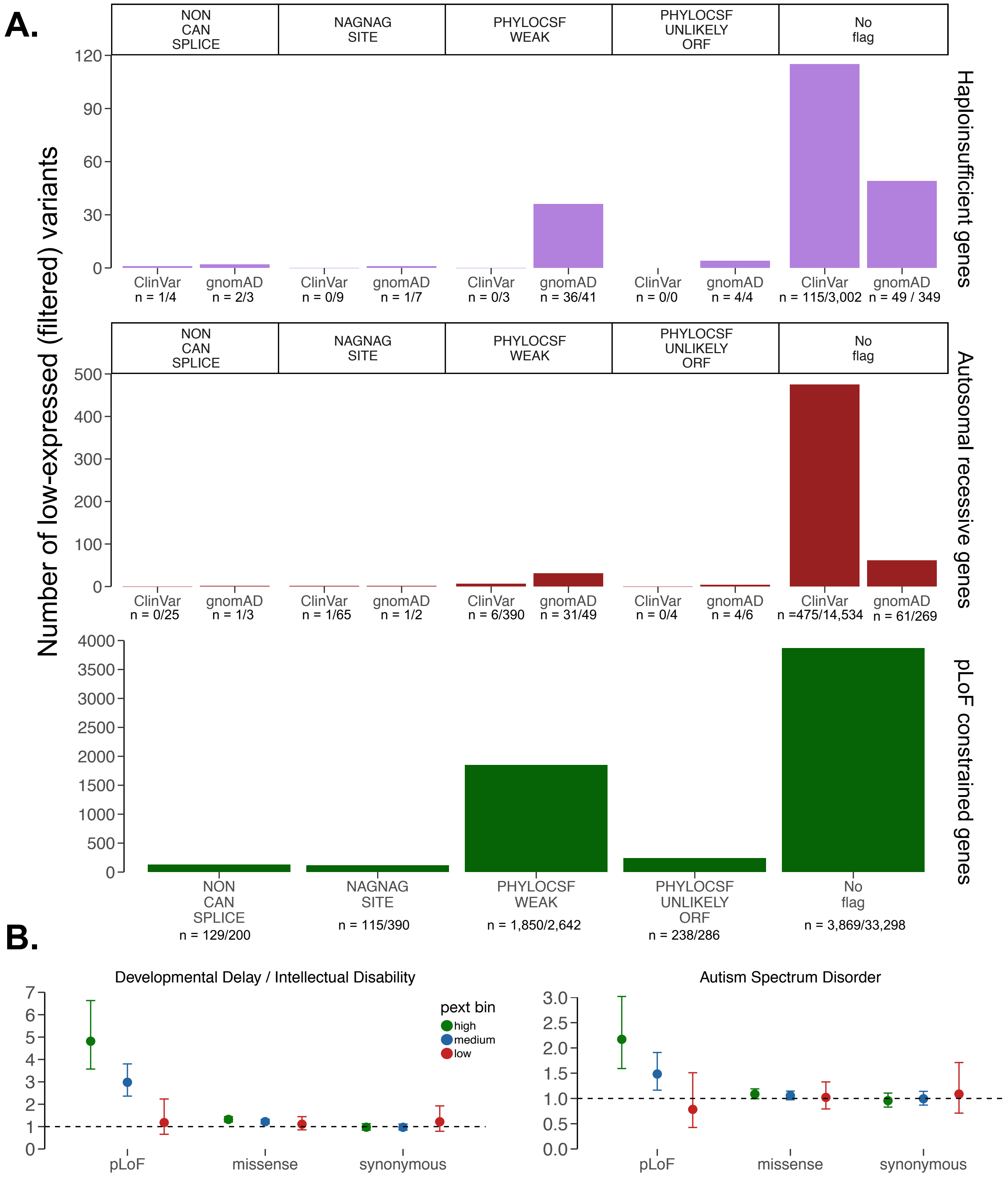


**Supplementary Figure 7: Evaluation of pext after accounting for LOFTEE flags in ClinVar and *de novo* analyses A.** Majority of pLoF variants filtered by pext are LOFTEE–HC with no flags. Comparison of proportion of high quality pLoF variants filtered in a curated list of 61 haploinsufficient developmental delays genes in gnomAD vs ClinVar (top), pLoF variants found in gnomAD in a homozygous state in at least one individual (middle) and pLoF and variants in gnomAD pLoF-intolerant genes (defined by LOEF < 0.35). Numbers below each x axis label represent the number of filtered variants over the number of total variants in the category. Synonymous variants are not shown for the bottom graph, given that synonymous variants do not receive LOFTEE designations. **B.** To show that results for *de novo* variant analysis do not change after controlling from LOFTEE flags, we regenerated Figure 4 restricted to only LOFTEE HC variants with no flags, removing 99 of all pLoF variants from a total of 2,161 (representing 4.6% of the analyzed pLoF variants).

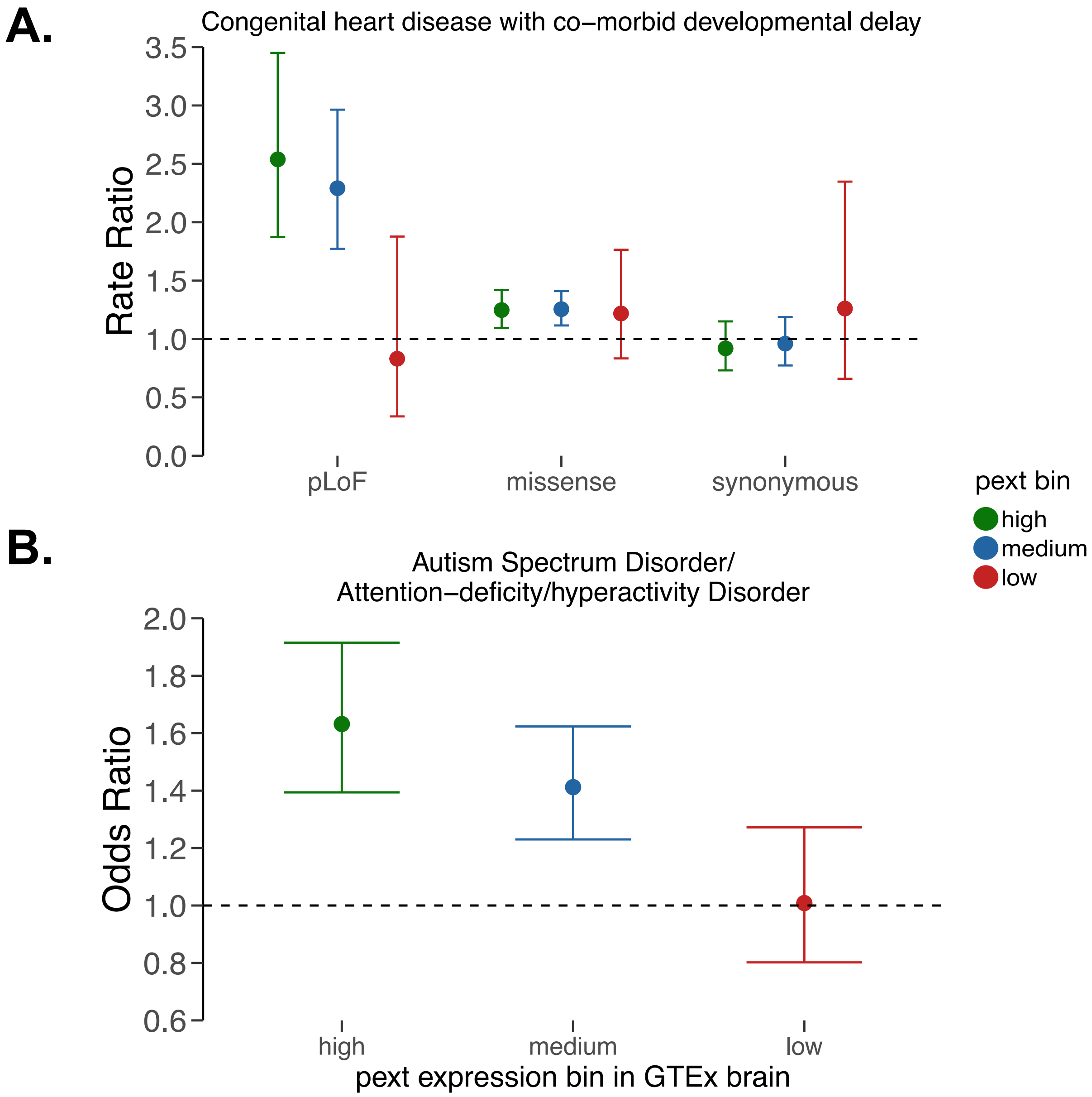


**Supplementary Figure 8: Application of transcript-expression based annotation to *de novo* and rare variant analysis in additional datasets A.** Using *de novo* variants identified in probands with congenital heart disease and co-morbid developmental delay we find a consistent effect of *de novo* pLoF variants found on high expressed regions in GTEx brain having larger effect sizes than *de novo* pLoF variants in weakly expressed regions. Once again, *de novo* pLoF variants found on regions with little evidence for expression are similarly distributed in cases and controls as *de novo* synonymous variants, suggesting such variants can be removed from gene burden testing analyses to boost discovery power. Rate ratio represents estimate from the poisson test. **B.** Rare pLoF variants (combined AC in cases and controls ≤ 10) identified in highly constrained genes (first decile in LOEUF) portioned upon pext expression bins show that those with high expression in GTEx brain have higher effect sizes than those identified in low-expressed regions, which are equally distributed in cases and controls. Odds ratio represents estimate from Fisher's exact text on counts.

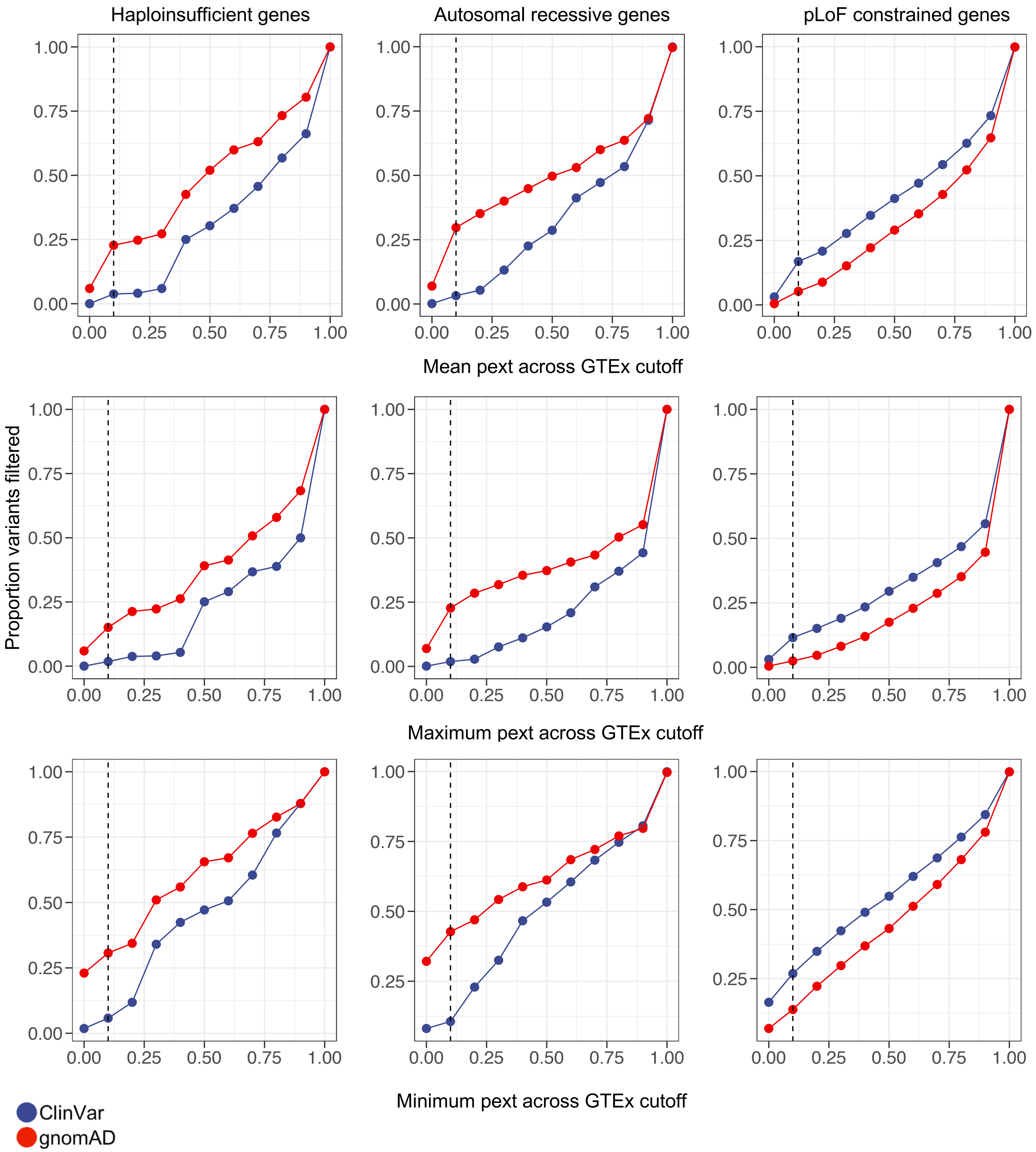


**Supplementary Figure 9: Evaluation of using differing cutoffs for average, minimum or maximum pext across tissues** While in this manuscript we have focused on binning average pext across GTEx tissues into low, medium and high-expressing categories, we also evaluated other summarizations of the GTEx pext scores in terms of the proportion of variants pLoF variants filtered from ClinVar and gnomAD in haploinsufficient severe disease genes, autosomal recessive disease genes, and pLoF constrained genes. All gene lists and variants represented are the same as used in Figure 4. We find that general trends of selective filtering of variants from gnomAD over ClinVar with pext is consistent across the varying methods for summarizing pext (average, minimum or maximum). Increasing the pext cut-off results in filtering more variants, and the given threshold used will depend on analysis needs. The dashed vertical line represents the 0.1 cutoff representing the low expression bin used for filtering in Figure 4.

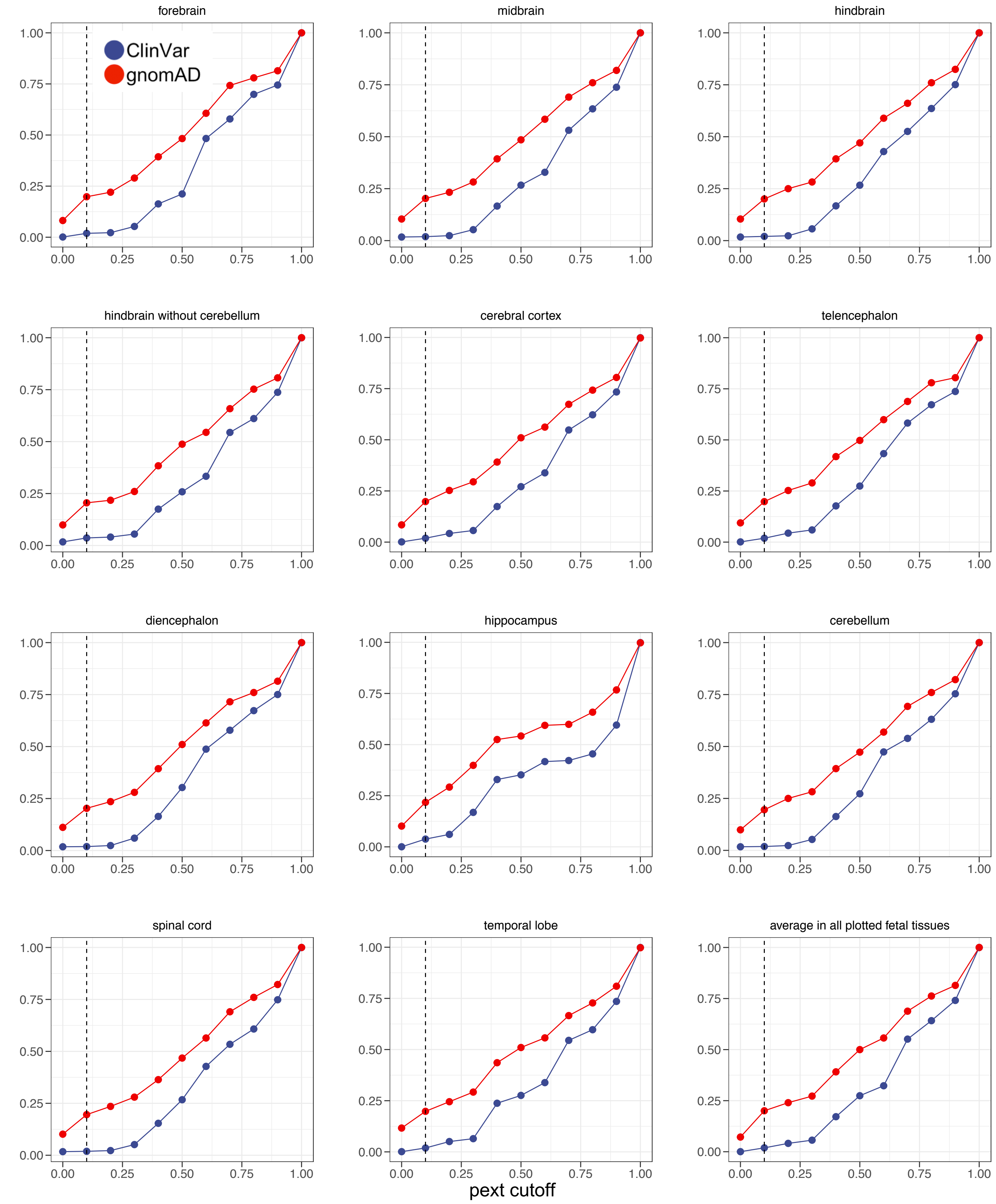


**Supplementary Figure 10: Using pext based on fetal RNA-seq isoform expression data for filtering variants in haploinsufficient genes** Given that the 61 haploinsufficient genes are likely expressed during development, we evaluated the proportion of variants filtered from gnomAD and ClinVar in these genes using pext scores derived from the Human Brain Developmental Resource (see methods). We observe that fetal-expression based pext does not provide more selective filtering than using GTEx, highlighting that pext-based filtering captures mostly annotation errors.

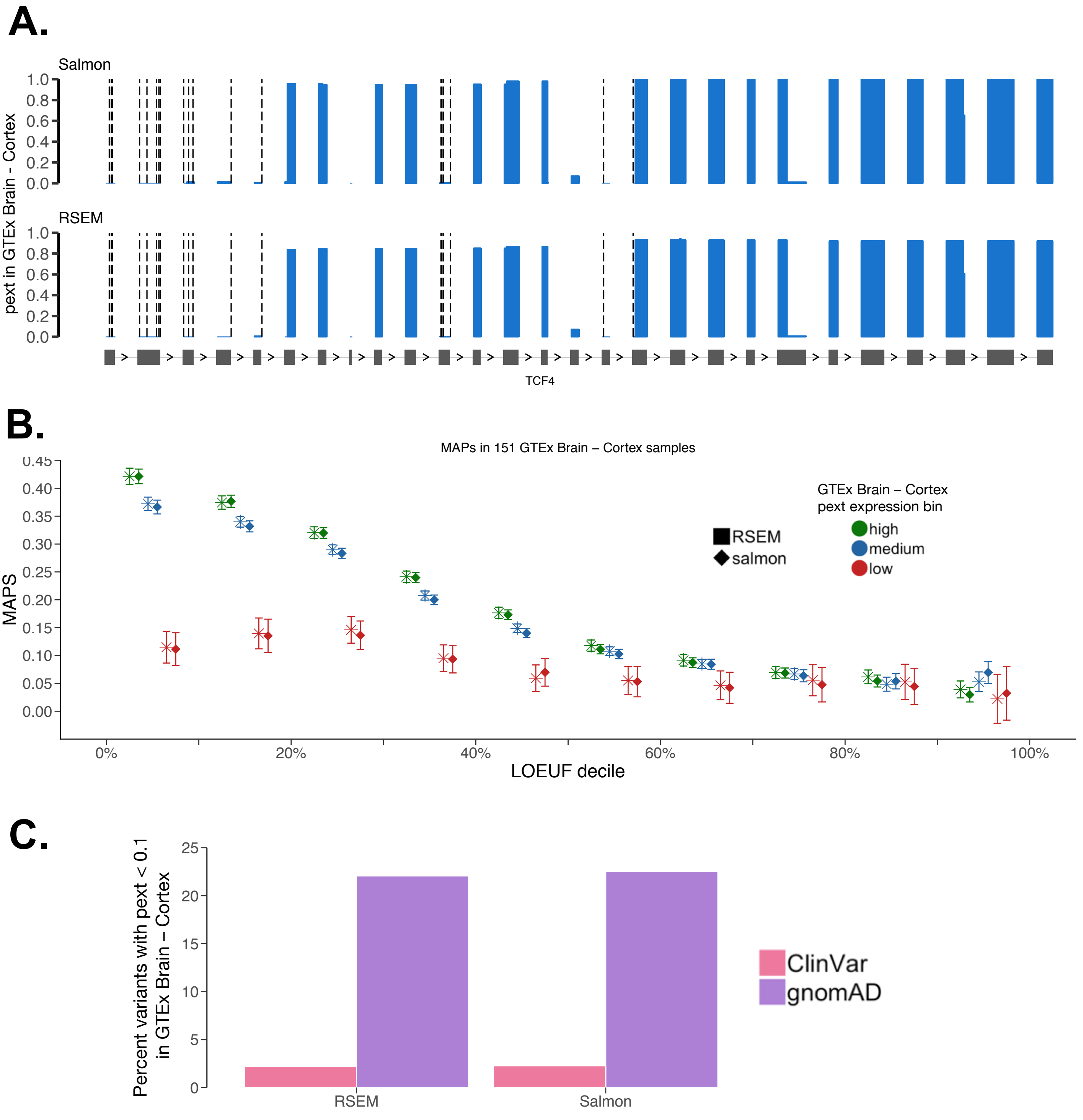


**Supplementary Figure 11 : Comparison of key results using Salmon vs RSEM.** Using isoform quantifications on 151 GTEx Brain – Cortex samples, we compared the results from analyses in the manuscript using quantifications from salmon and RSEM. Results from the analyses were consistent, suggesting that the pext calculation is robust to isoform quantification tool used. To keep analysis consistent, genes where the maximum pext was below 0.2 were removed, resulting in the filtering of 691 and 325 genes from the RSEM and salmon quantifications, respectively (see Methods) **A.** The baselevel pext metric in *TCF4* using the two quantification tools. The 20 pLoF variants identified in gnomAD in *TCF4*, denoted by dashed lines, lie on unexpressed regions in Brain – Cortex samples using salmon and RSEM **B.** No significant difference in the MAPs score is seen for pLoF variants in pext expression bins with RSEM and salmon quantifications. **C.** The number of pLoF variants filtered with a Brain – Cortex pext cutoff of 0.1 in gnomAD vs ClinVar was similar.

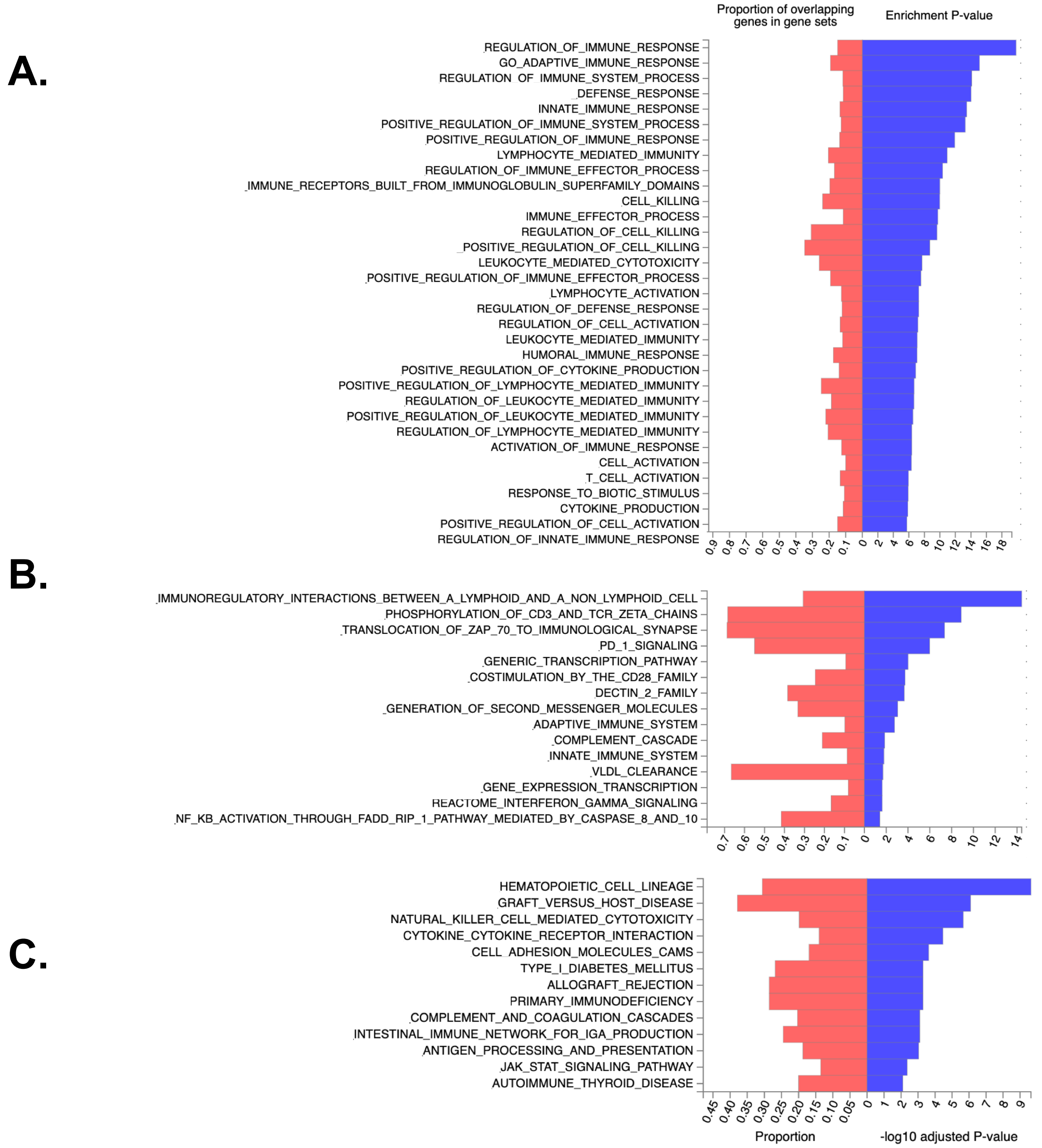


**Supplementary Figure 12: Results from FUMA GENE2FUNC analysis in unconserved regions with high expression values** We ran pathway analysis on 1,308 genes harboring 2,411 regions with low conservation (phyloCSF < -100) but high expression (pext > 0.9) shown in Figure 3A using the FUMA GENE2FUNC web browser. Results from **A)** Gene Ontology Biological Processes **B)** Reactome pathways and **C)** KEGG pathways show that these regions are enriched for immune pathways, which are selected for diversity but represent true coding regions, emphasizing the orthogonal information provided by pext over conservation alone. Full results from FUMA analysis are available in Supplementary Table 5

### Supplementary Table 1: Summary of manual curation flags for 401 pLoF in 61 haploinsufficient disease genes identified in gnomAD

| Mapping error | Strand Bias | Reference error | Genotyping error | Homopolymer | MNV | Essential splice site rescue | Minority of transcripts | Weak exon conservation | Last exon | Other annotation error |
|---|---|---|---|---|---|---|---|---|---|---|
| Human self-chained repeats (UCSC) | Variant present on >90% of either forward or reverse strand | Reference contains small (<5bp) deletion within exon | Allele balance ≤ 35% | Repeat of ≥ 6 base pairs | In phase MNV that abolishes stop codon | In frame splice site rescue within 36 bp and validated by Alamut | Variant falls on <50% of coding NCBI RefSeq transcripts | Flagged by phyloCSF as weak | Variant falls within the terminal coding exon | Variant falls on exactly 50% of NCBI coding RefSeq transcripts |
| Tandem repeats (UCSC) | | | GC rich region | Repeat of ≥ 6 trinucleotide repeats | Frame restoring indel | | | Entire exon is weakly conserved upon visual inspection in the UCSC browser | Variant falls within 50 base pairs of penultimate coding exon | Exon is partially conserved, specifically lacking conservation of variant/transcript |
| Segmental dups (UCSC) | | | GQ ≤ 20 | | | | | | Variants affects >25% coding sequence | Re-initiation by downstream methionine in first coding exon |
| Complex variation in region e.g. multiple indels, SNVs | | | Low complexity sequence | | | | | | | Stop codon occurs upstream of variant |
| | | | Low read depth <15 | | | | | | | |

### Supplementary Table 2: Summary of criteria for LoF verdicts of 401 pLoF in 61 haploinsufficient disease genes identified in gnomAD

| LoF | Likely LoF | Likely not LoF | Not LoF |
|---|---|---|---|
| GQ > 99, absence of any evidence to refute a LOF consequence and one of two criteria met: AB>35 and read depth >15 | Low complexity region | Re-initiation by downstream methionine in first coding exon | Minority of coding RefSeq transcripts (except when exon well conserved) |
| | Allele balance ≤ 35% but >25% | Mapping ambiguity (UCSC) | Variant falls in last coding exon |
| | GQ ≤ 20 | phyloCSF weak | Weak conservation of exon |
| | GC rich region | In frame splice site rescue between 6 and 21 base pairs from the intron/exon boundary and validated by Alamut | Frame restoring indel |
| | Strand bias in regions where coverage is skewed towards a strand | Homopolymer repeat | In phase multi-nucleotide variant abolishing stop codon |
| | Read depth <15 | Falls within terminal exon although will disrupt >25% but <50% of coding sequence | Complete splice site rescue within 6 base pairs and validated in Alamut |
| | Any single other transcript error | Minority of transcripts but exon well conserved | Complex mapping and assembly error |
| | Potential splice site rescue >21 BP away and weakly supported by Alamut | Combination of multiple flags (e.g. On 50% of RefSeq transcripts and partial loss of exon conservation; homopolymer repeat and mapping error etc.) | Reference error |
| | Splice rescue within 21 BP but very weak signal as per Alamut | Strand bias despite equal coverage of forward and reverse strand across region | Combination of multiple flags (e.g. in frame splice rescue within 21 bp and minority of transcripts with exon well conserved; mapping ambiguity and phyloCSF weak etc.) |
| | Falls within terminal exon although will disrupt >50% of coding sequence | | |
