## Supplementary material for "Transcript expression-aware annotation improves rare variant discovery and interpretation": upplementary Figure 4 - Baselevel TCF4 expression per GTEx tissue

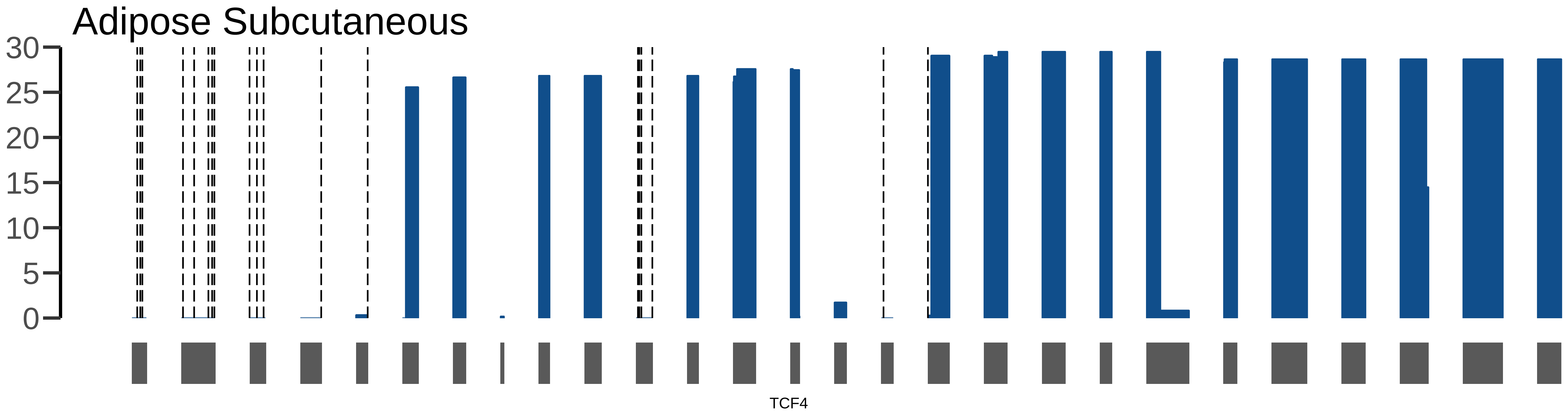

Adipose Subcutaneous
30
25
20
15
10
5
0
TCF4

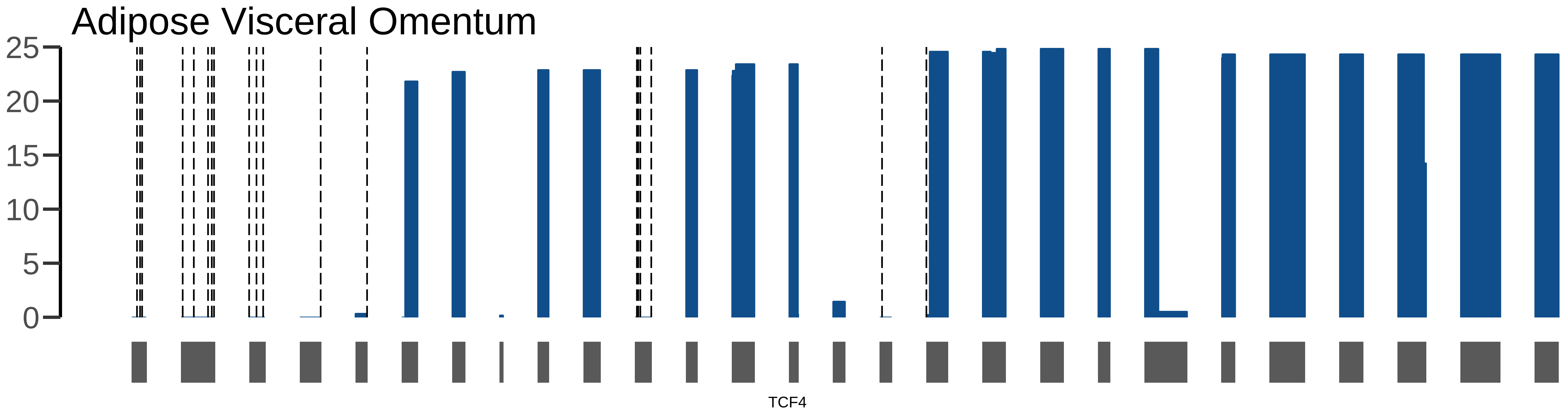

Adipose Visceral Omentum
25
20
15
10
5
0
TCF4

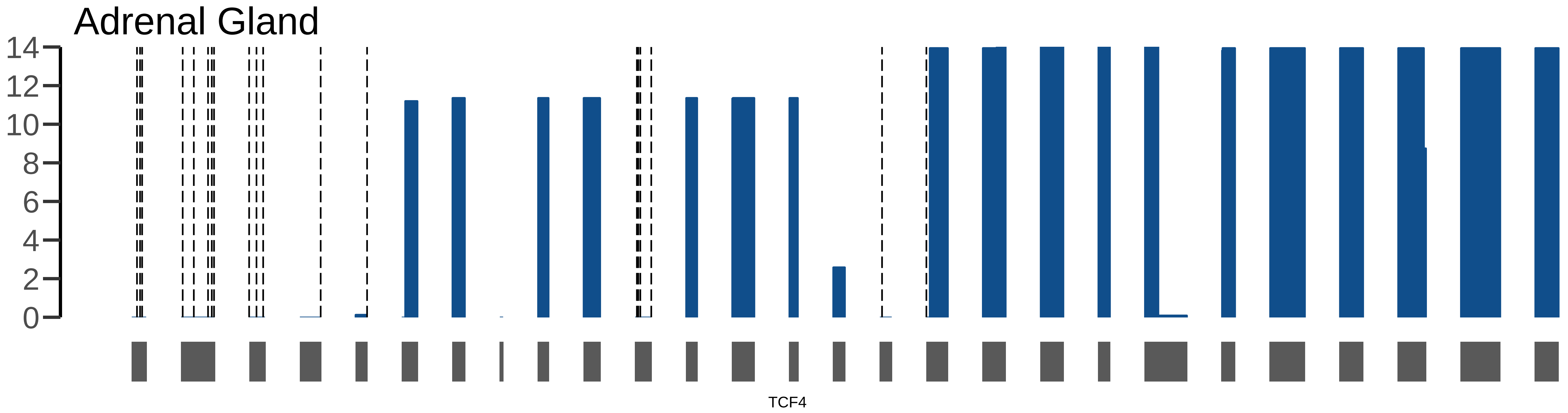
Adrenal Gland
14
12
10
8
6
4
2
0
TCF4

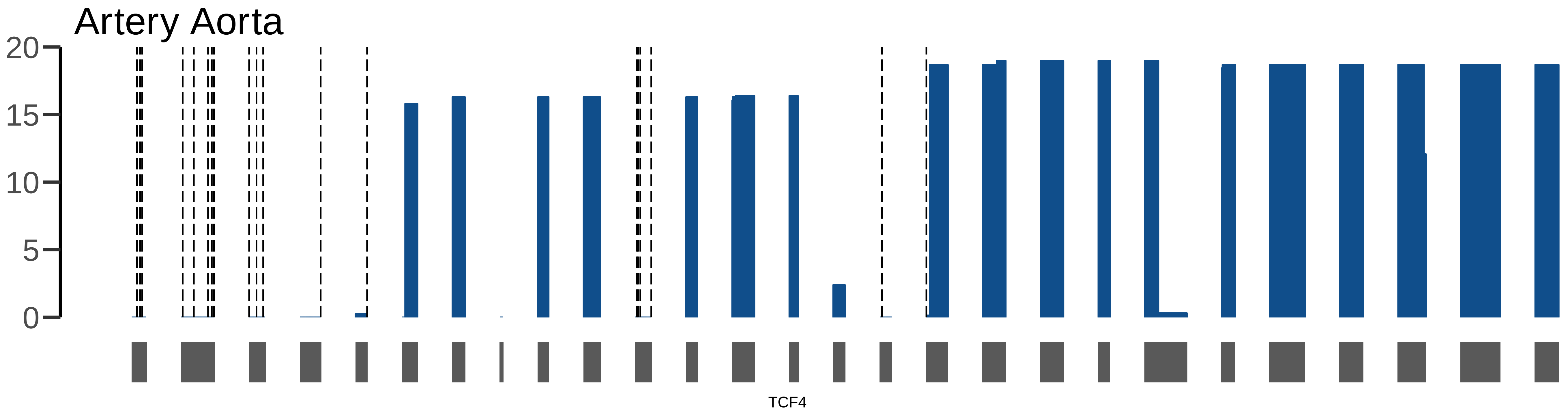

Artery Aorta
20
15
10
5
0
TCF4

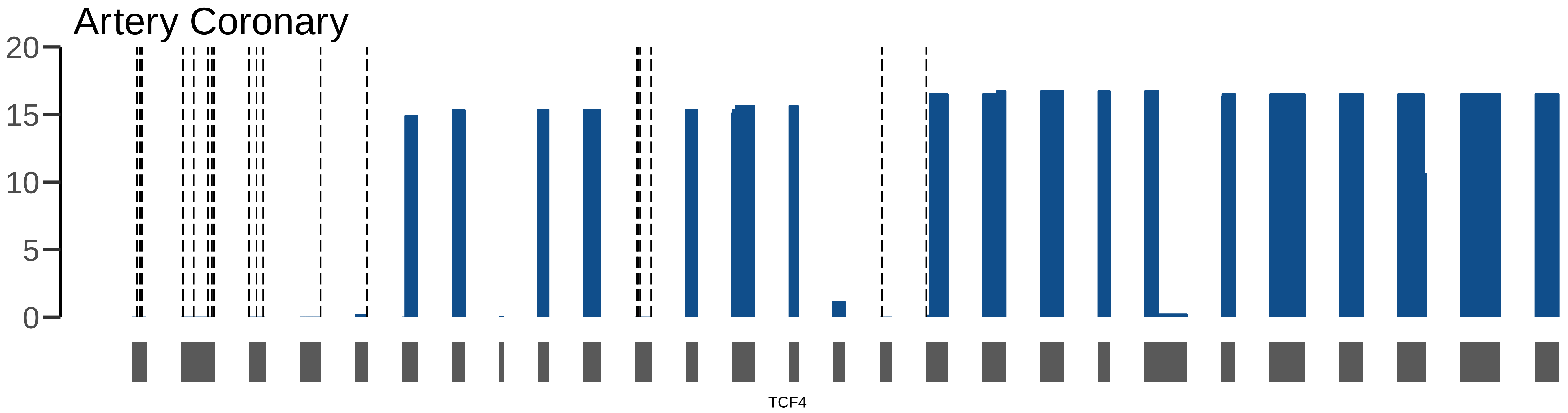

Artery Coronary
20
15
10
5
0
TCF4

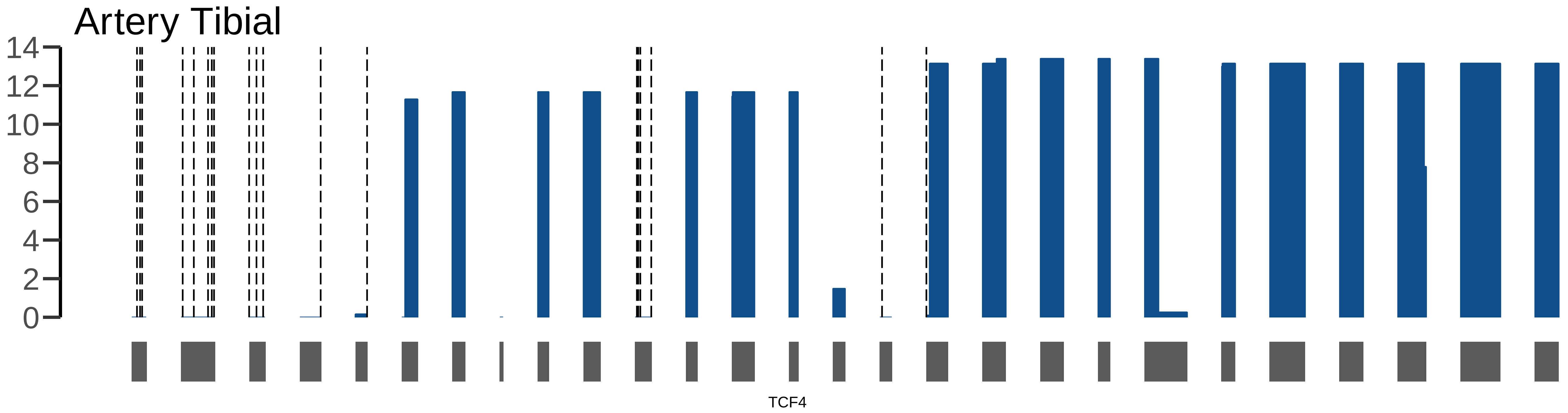
Artery Tibial
14
12
10
8
6
4
2
0
TCF4

#### Brain Amygdala

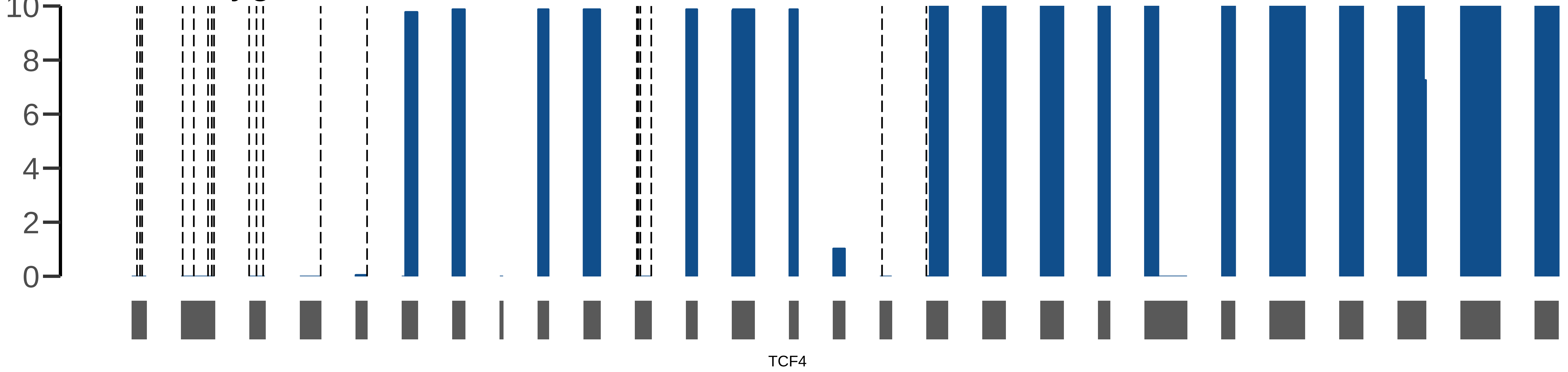

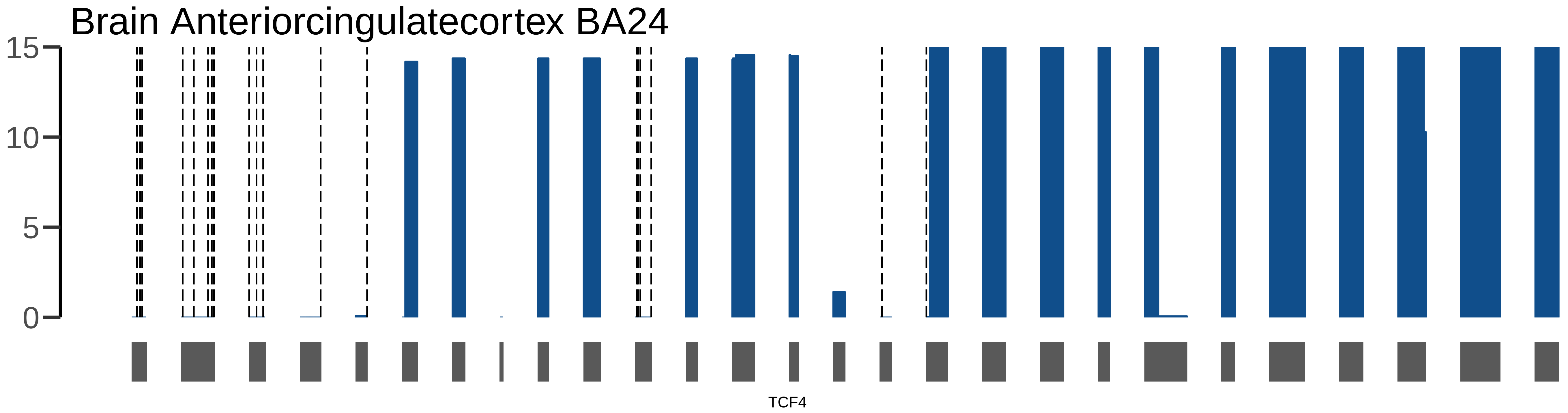
Brain Anteriorcingulatecortex BA24
15
10
5
0
TCF4

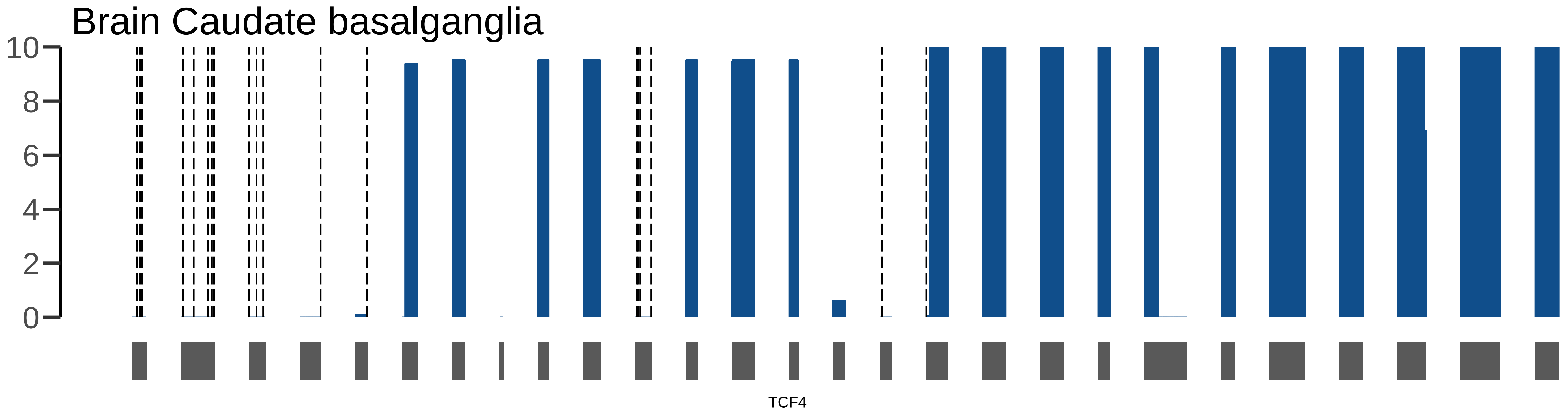

Brain Caudate basalganglia
10
8
6
4
2
0
TCF4

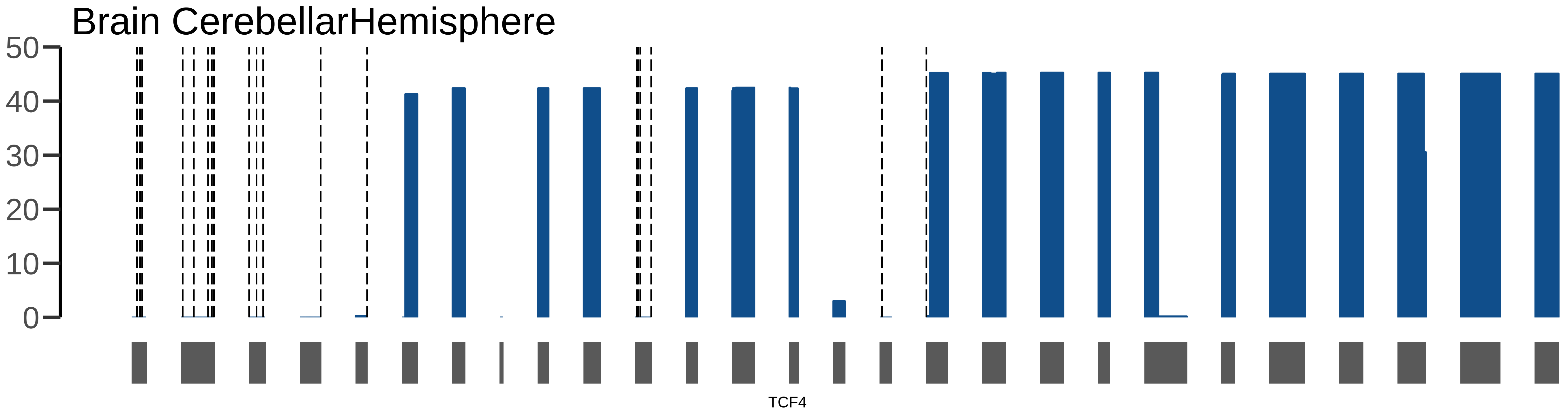
Brain CerebellarHemisphere
50
40
30
20
10
0
TCF4

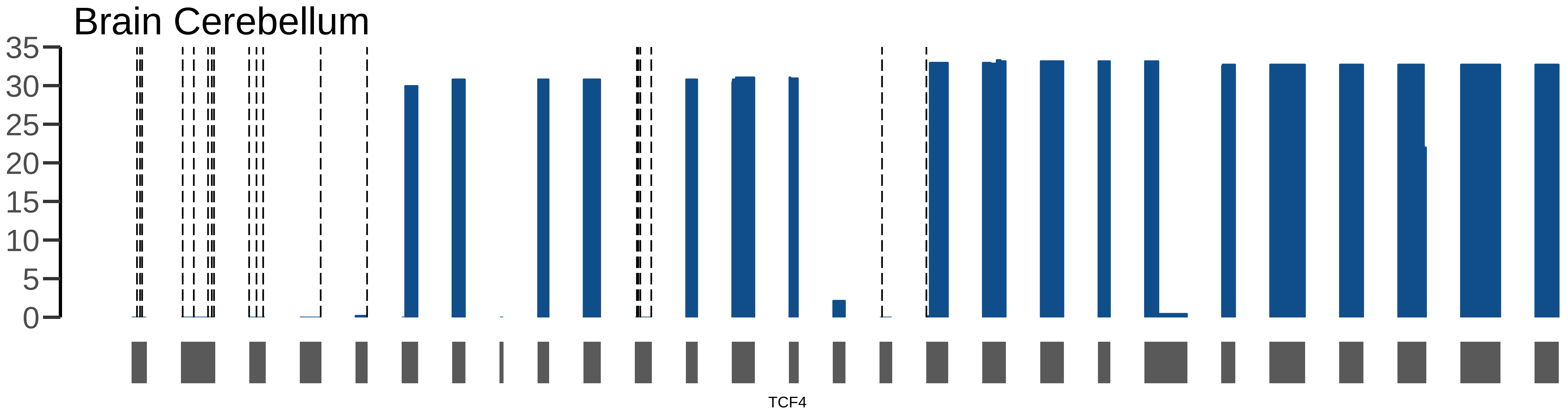
Brain Cerebellum
35
30
25
20
15
10
5
0
TCF4

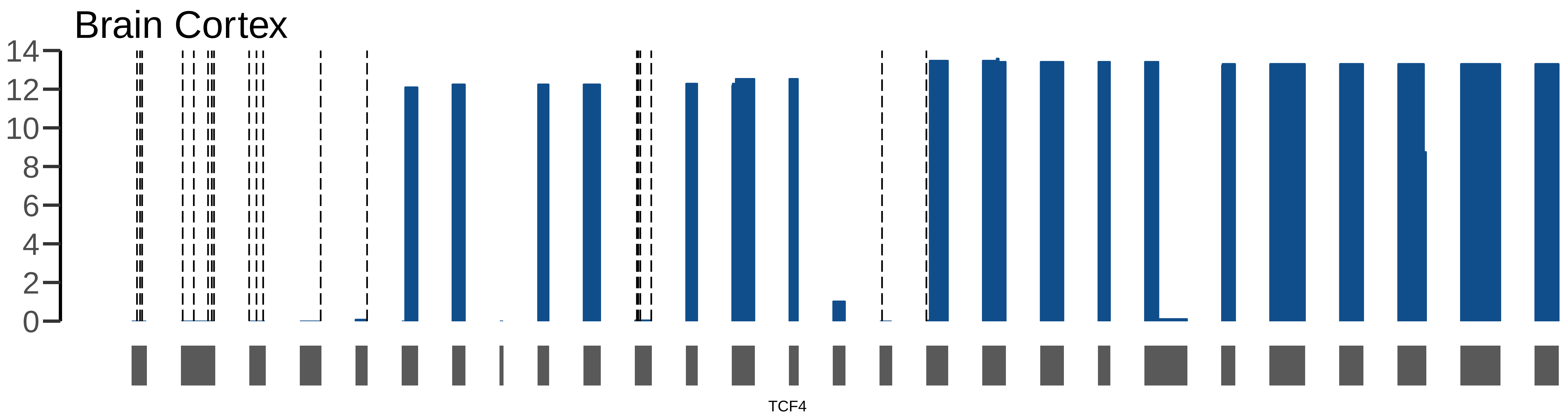
Brain Cortex
14
12
10
8
6
4
2
0
TCF4

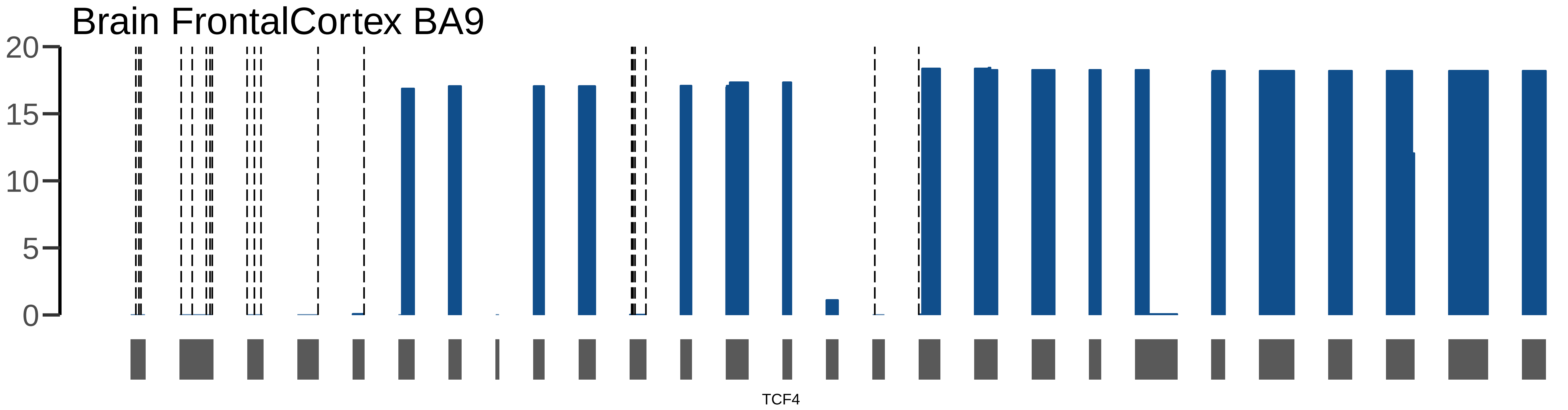
Brain FrontalCortex BA9
20
15
10
5
0
TCF4

### Brain Hippocampus

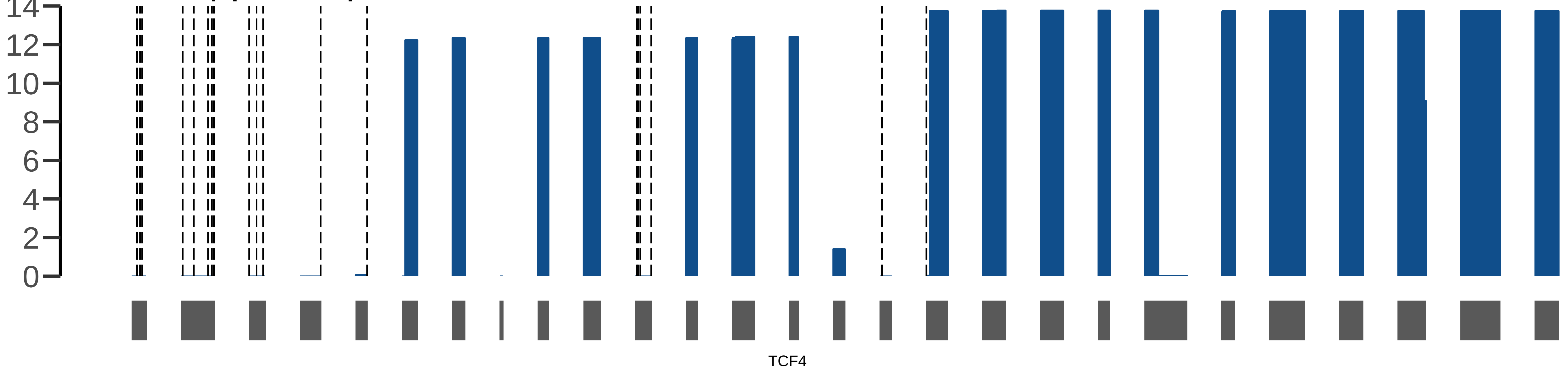

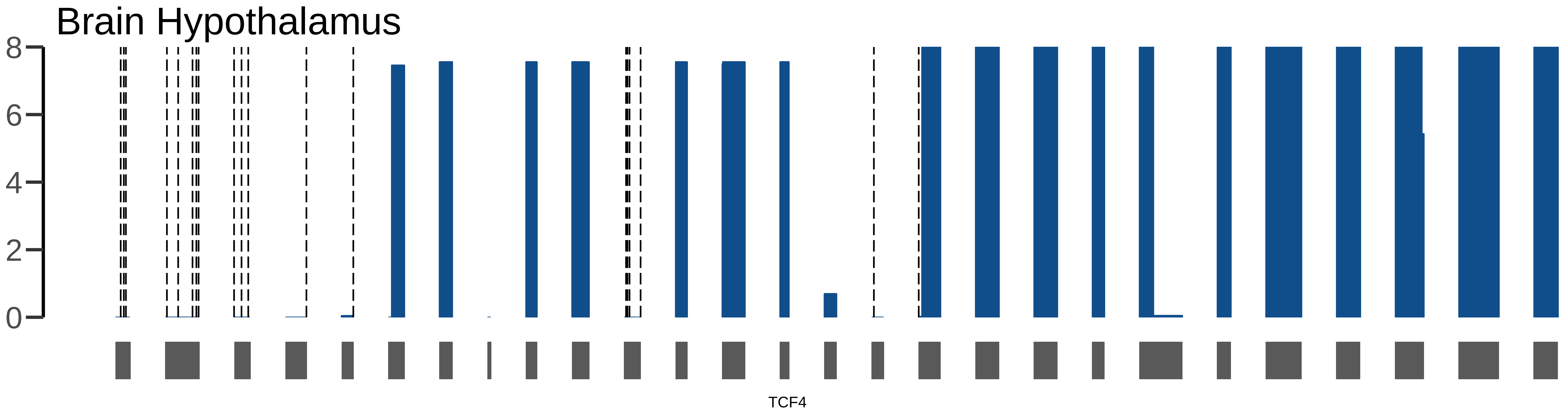
Brain Hypothalamus
8
6
4
2
0
TCF4

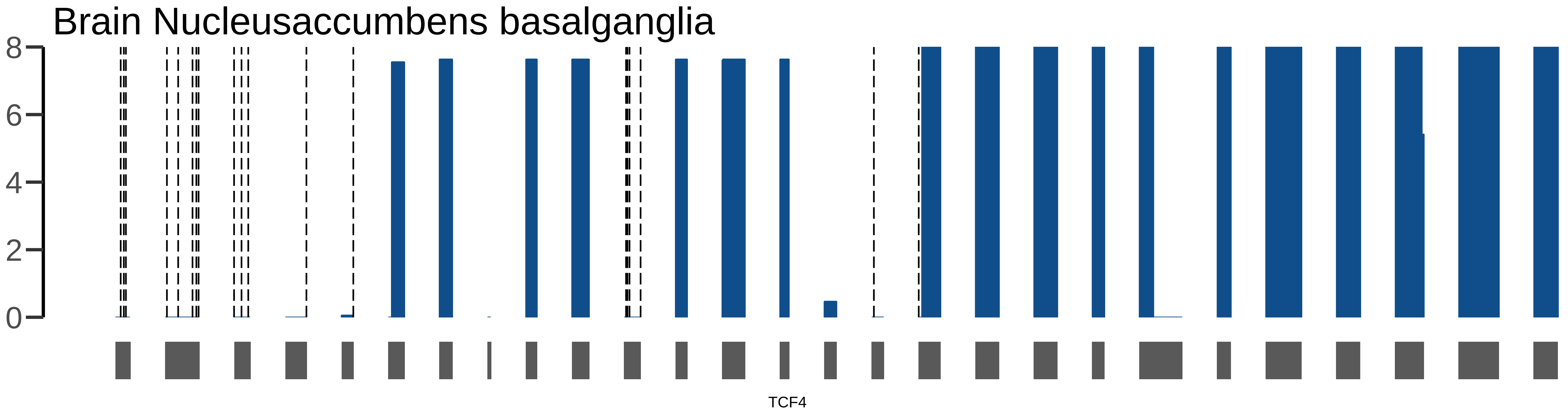

Brain Nucleusaccumbens basalganglia
8
6
4
2
0
TCF4

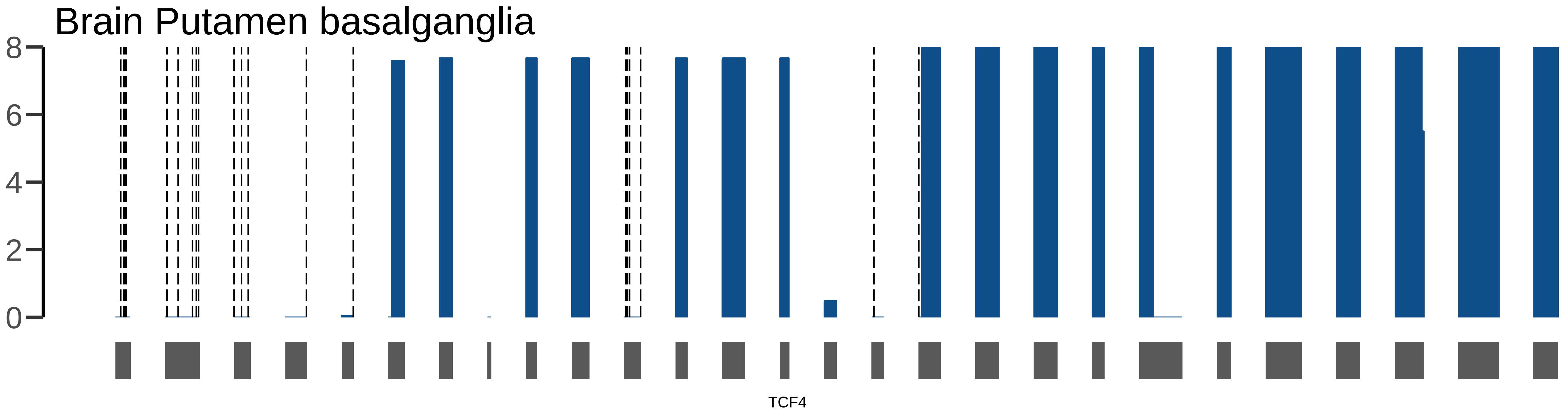
Brain Putamen basalganglia
8
6
4
2
0
TCF4

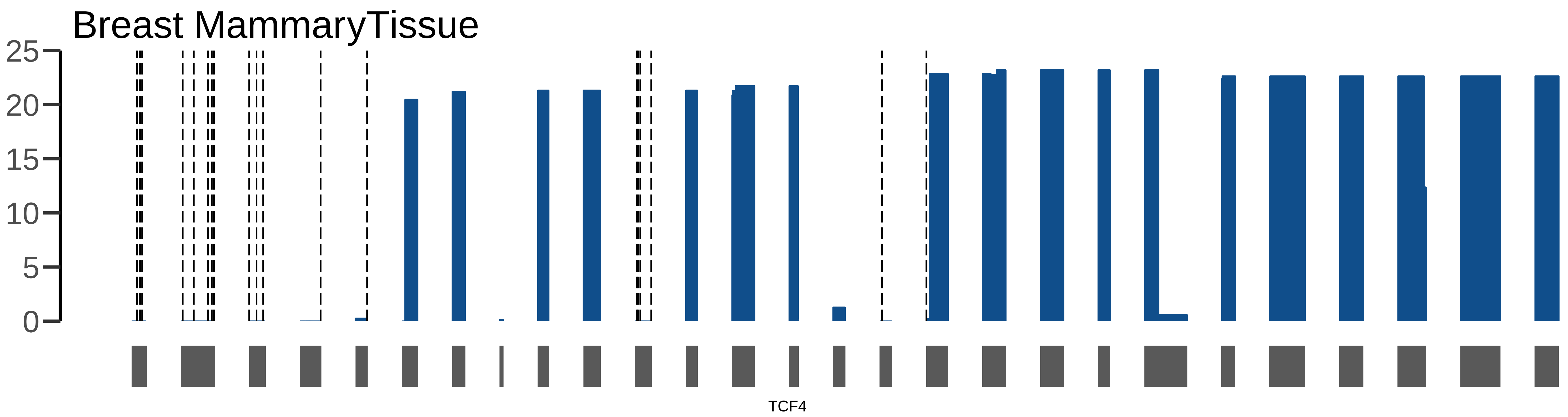

Breast MammaryTissue
25
20
15
10
5
0
TCF4

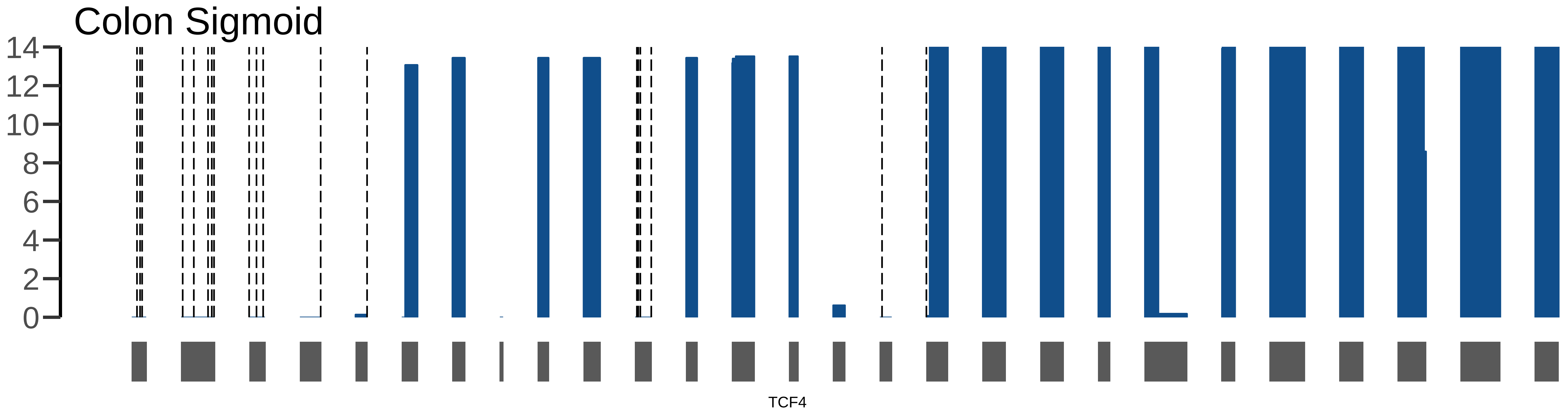

Colon Sigmoid
14
12
10
8
6
4
2
0
TCF4

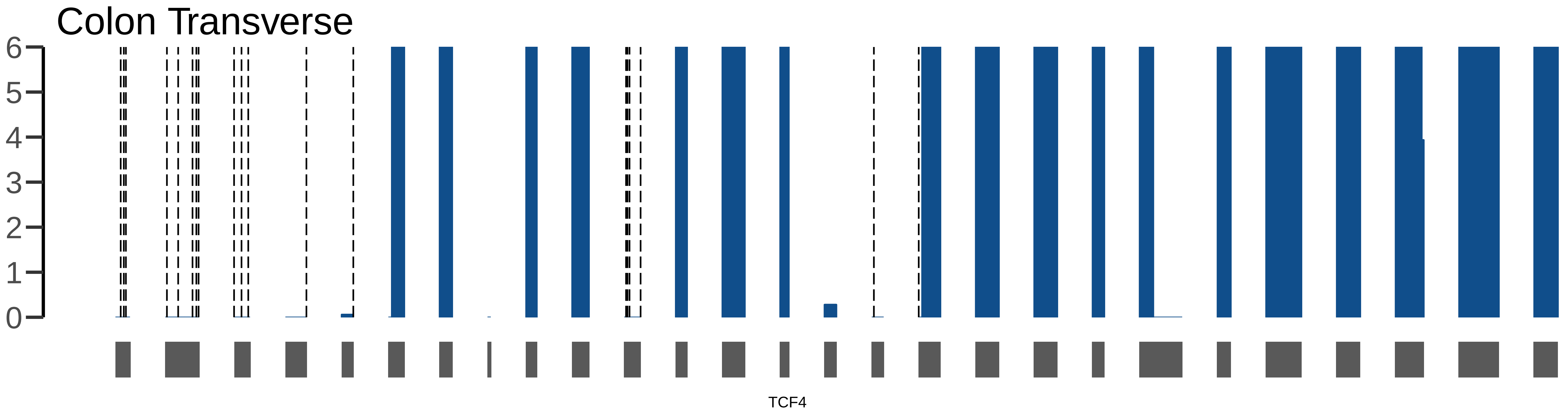

Colon Transverse
6
5
4
3
2
1
0
TCF4

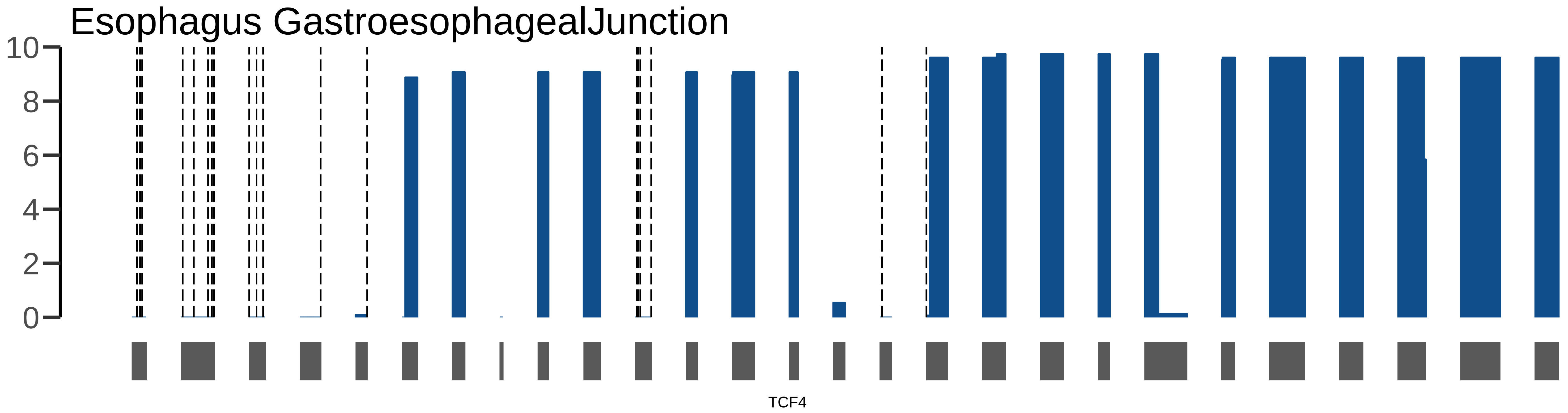

Esophagus GastroesophagealJunction
10
8
6
4
2
0
TCF4

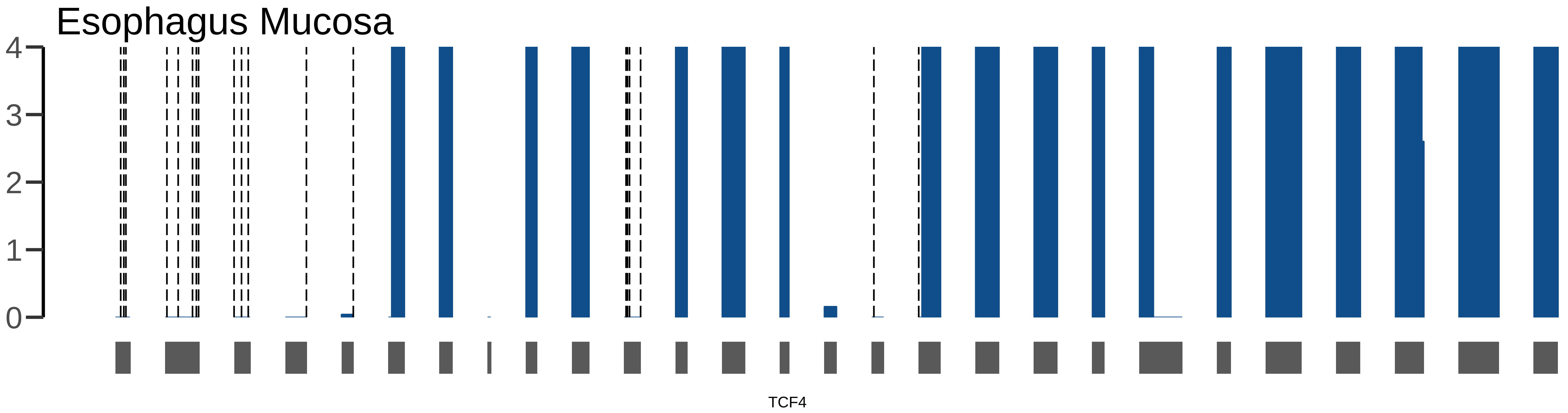
Esophagus Mucosa
4
3
2
1
0
TCF4

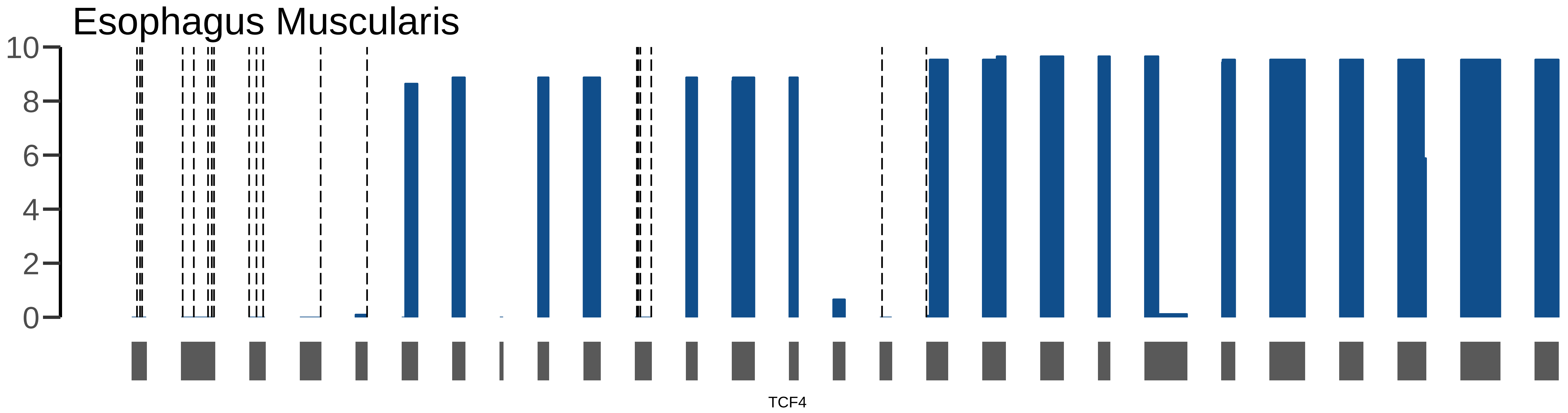
Esophagus Muscularis
10
8
6
4
2
0
TCF4

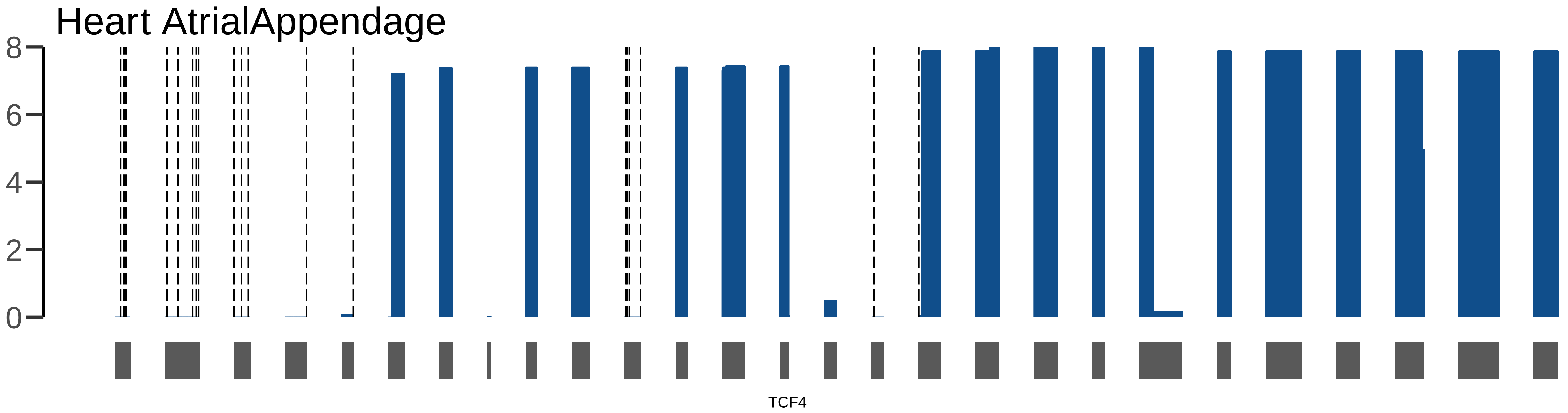
Heart AtrialAppendage
8
6
4
2
0
TCF4

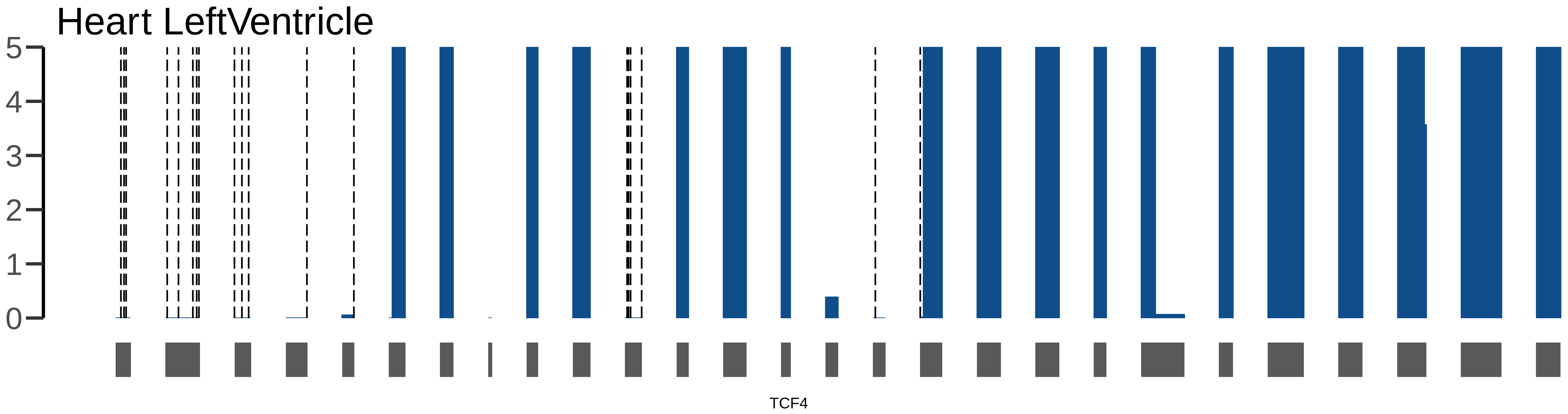
Heart LeftVentricle
5
4
3
2
1
0
TCF4

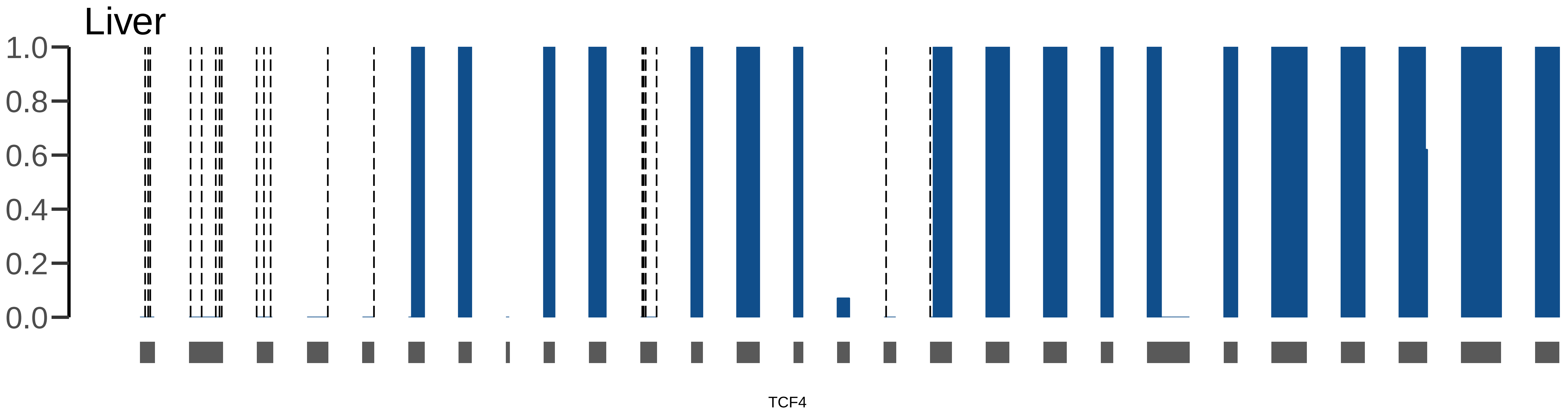

Liver
1.0
0.8
0.6
0.4
0.2
0.0
TCF4

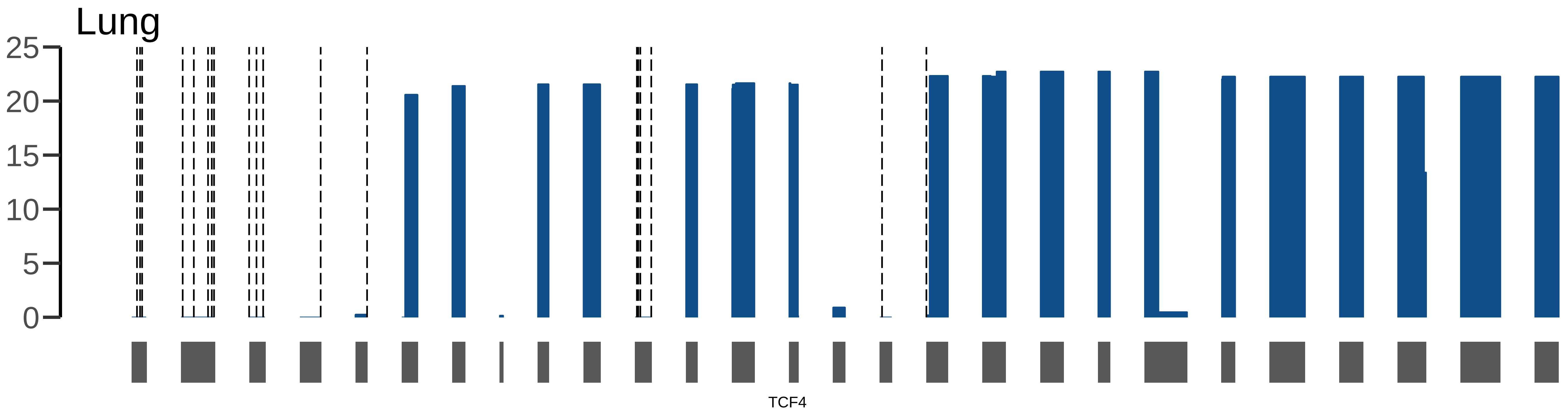
Lung
25
20
15
10
5
0
TCF4

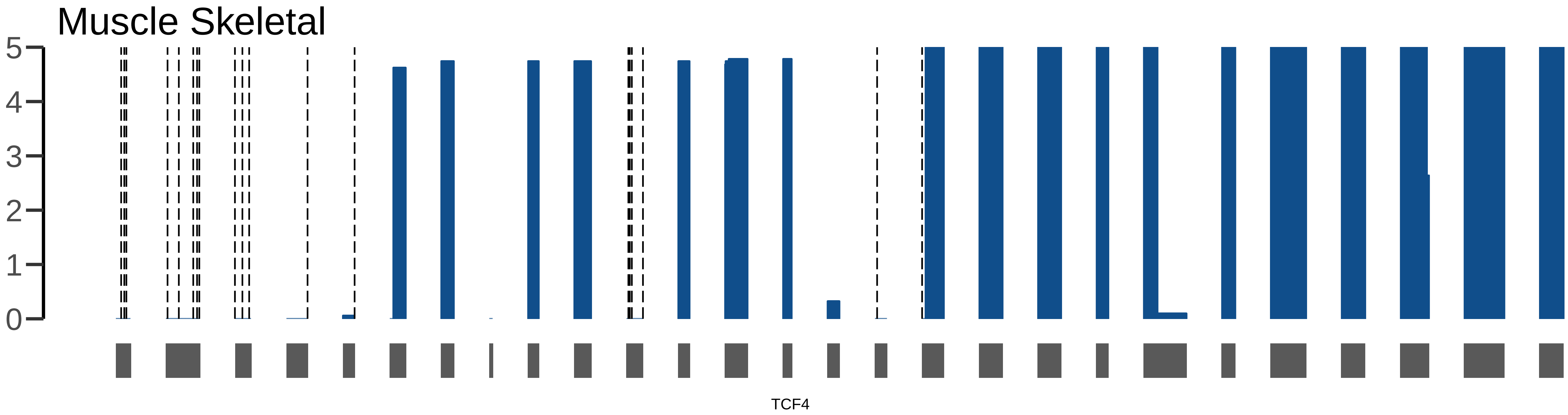
Muscle Skeletal
5
4
3
2
1
0
TCF4

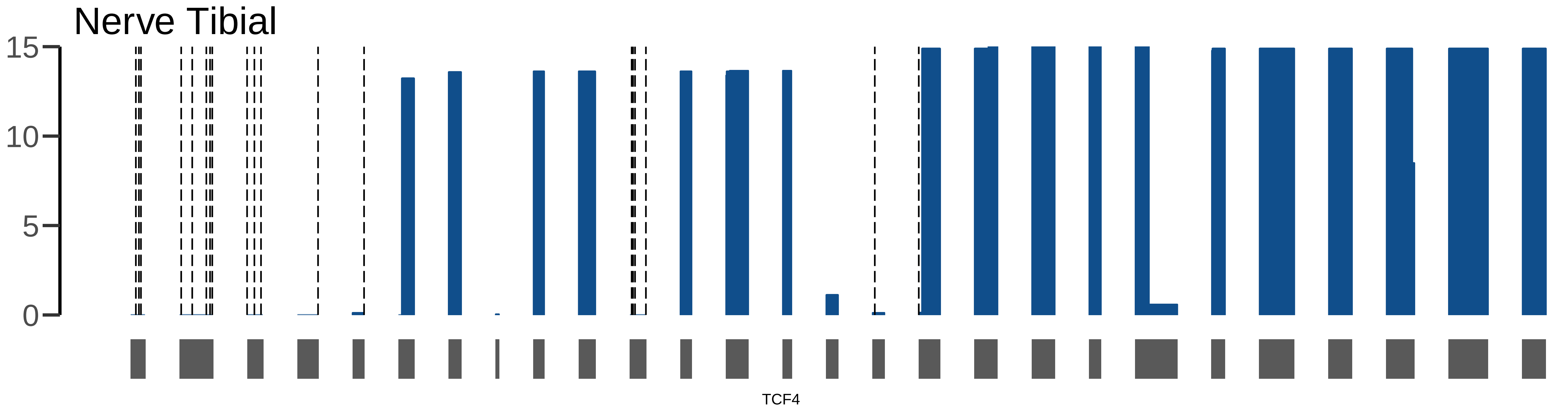
Nerve Tibial
15
10
5
0
TCF4

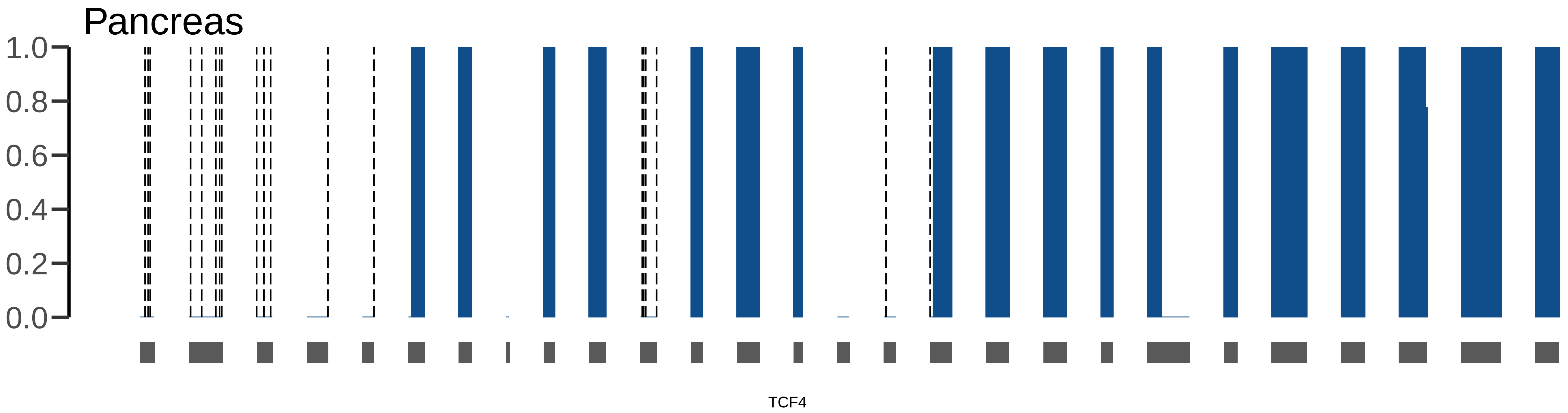
Pancreas
1.0
0.8
0.6
0.4
0.2
0.0
TCF4

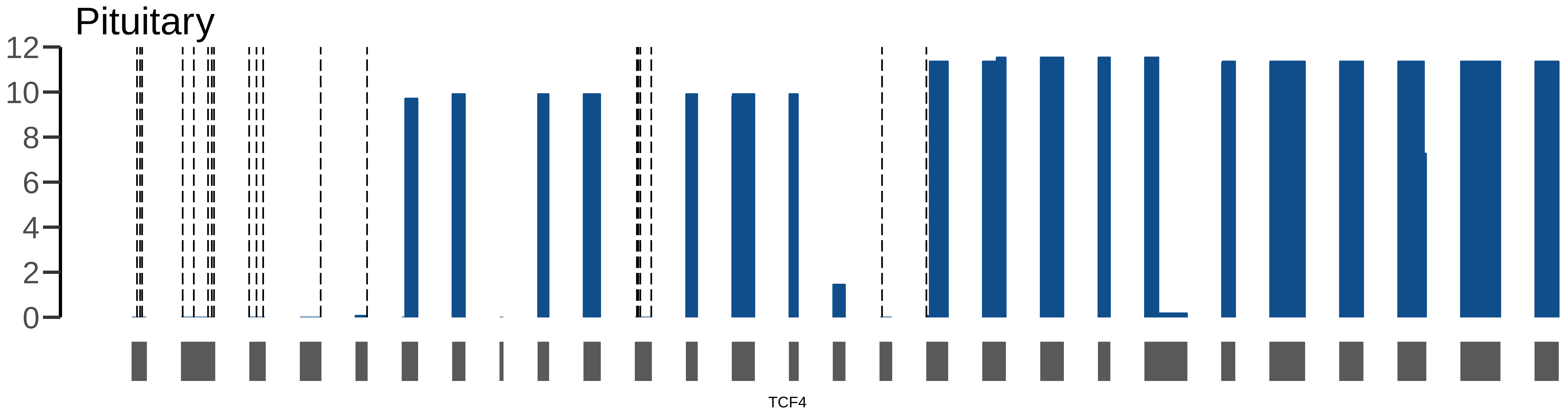

Pituitary
12
10
8
6
4
2
0
TCF4

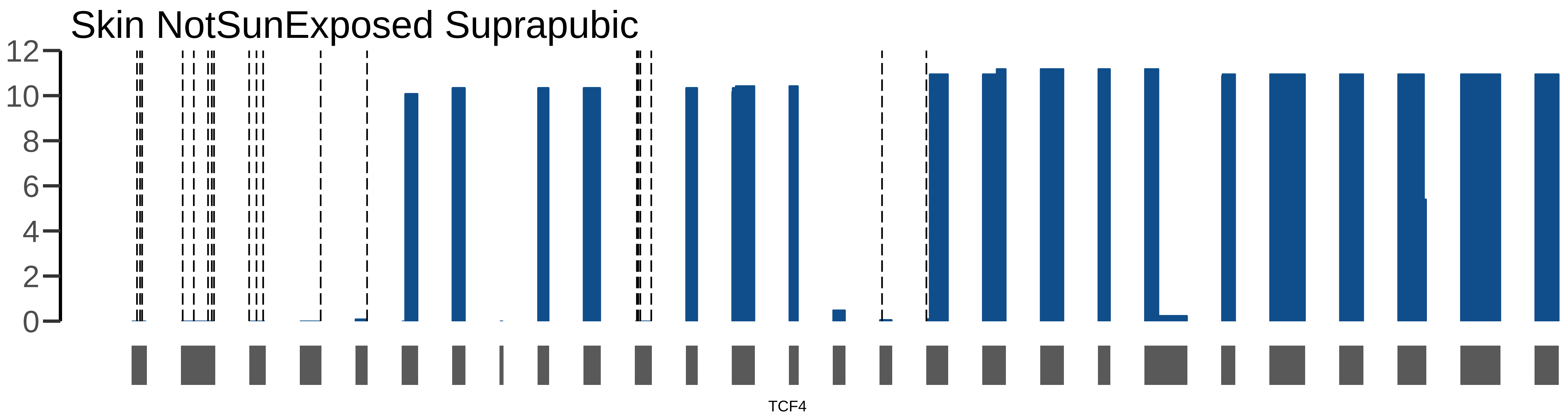
Skin NotSunExposed Suprapubic
12
10
8
6
4
2
0
TCF4

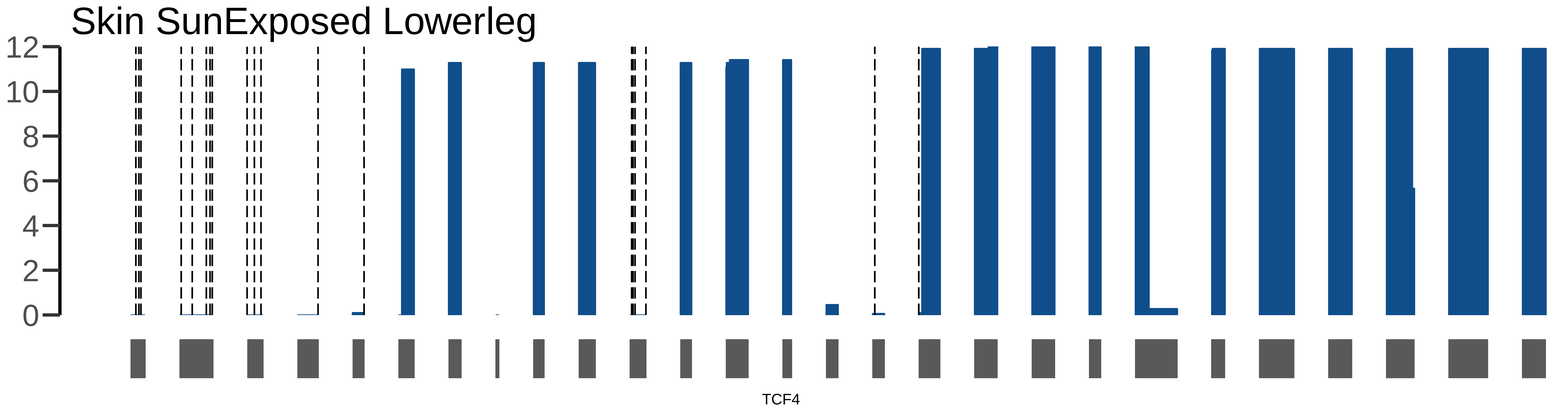
Skin SunExposed Lowerleg
12
10
8
6
4
2
0
TCF4

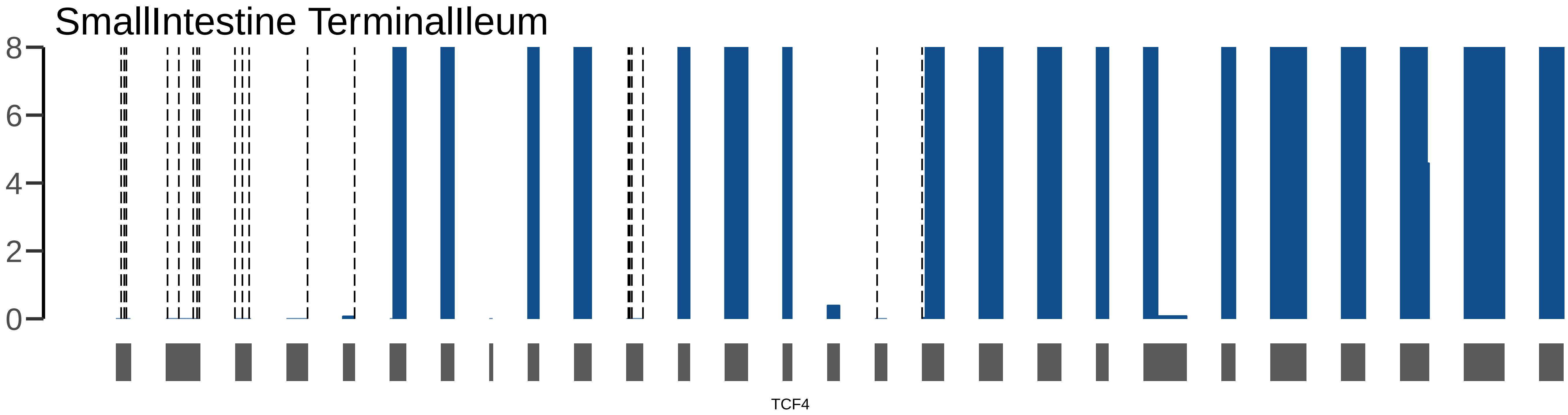
SmallIntestine TerminalIleum
8
6
4
2
0
TCF4

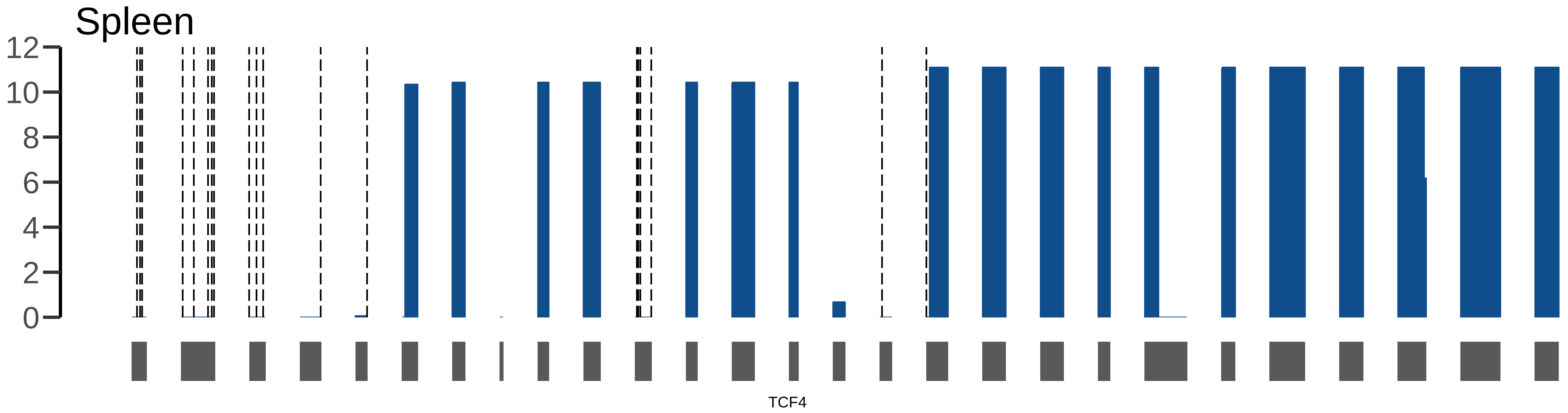
Spleen
12
10
8
6
4
2
0
TCF4

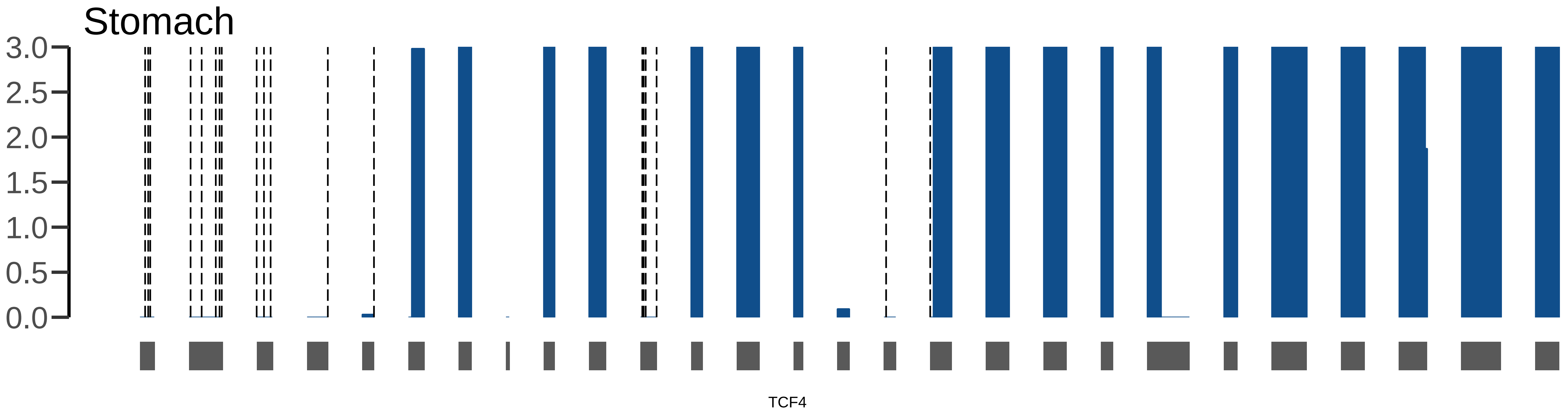
Stomach
3.0
2.5
2.0
1.5
1.0
0.5
0.0
TCF4

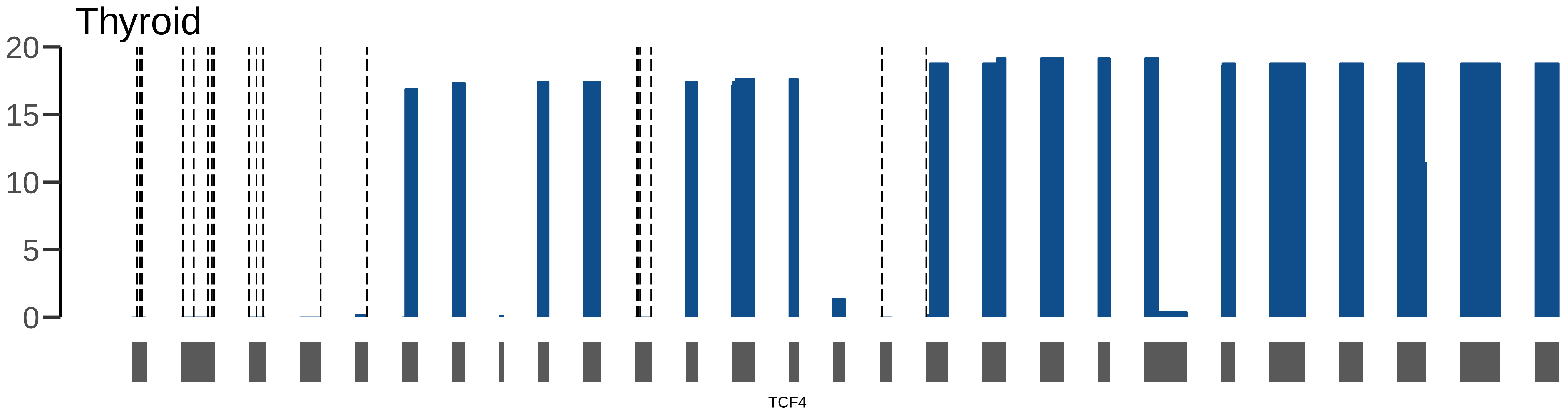

Thyroid
20
15
10
5
0
TCF4

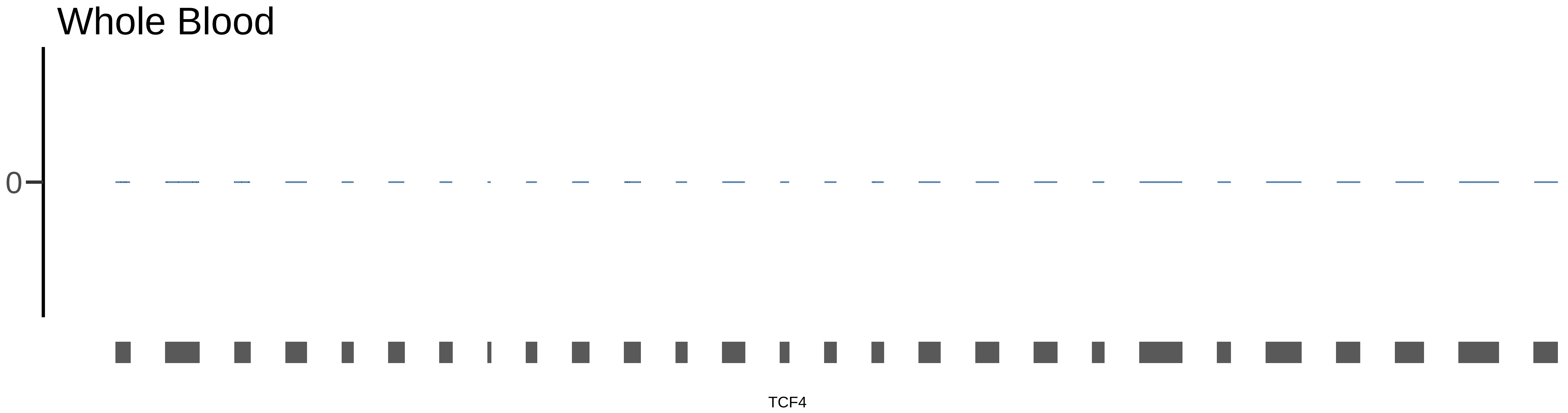

Whole Blood
0
TCF4
